# Phylogenetic Measures of the Core Microbiome

**DOI:** 10.1101/2024.10.10.617603

**Authors:** Sharon Anne Bewick, Benjamin Thomas Camper

## Abstract

**Background:** A useful concept in microbial ecology is the ‘core microbiome.’ Typically, core microbiomes are defined as the microbial taxa, genes, or functions shared by a threshold number of microbiome samples from a particular type of habitat (e.g., a particular type of host or a particular type of environment/ecosystem). In defining the core microbiome, the goal is to capture the portion of the microbial community that is conserved across samples from the focal habitat. Recently, there has been growing interest in developing methods to better characterize core microbiomes. As a result, numerous occurrence- and abundance-based measures have been defined. However, few have included phylogeny-aware metrics for analyzing core microbiomes.

**Results:** In this paper, we develop the concept of the ‘core community phylogeny’ – a phylogeny where branches are selected based on their presence in multiple samples from a single type of habitat. We then use the core community phylogeny to define phylogenetic metrics describing the diversity of core microbiomes from a single type of habitat, the turnover between core microbiomes from two different types of habitats, and the shared diversity across core microbiomes from two or more different types of habitats. As compared to non-phylogenetic metrics, our phylogenetic metrics show greater consistency across taxonomic rank and/or phylogenetic level, as well as less sensitivity to strain variation across microbiome samples. Thus, our metrics address key challenges in the interpretation of core microbiomes.

**Conclusions:** We provide a phylogenetic framework for characterizing and comparing core microbiomes. Importantly, the methods that we propose allow seamless integration of microbiome properties across taxonomic rank and/or phylogenetic level. Ultimately, this will provide both a more consistent picture of the core microbiome, as well as novel biological insight into the conserved components of microbial communities.

## Introduction

Core microbiomes have become increasingly important as a dimension for studying and characterizing sets of microbial communities.^1,2^ Broadly speaking, a core microbiome is the group of microbial taxa, genes, or functions shared by a large number of microbiome samples from a specific type of habitat. Often, the habitat is defined based on a host species^3–5^ (e.g., the core microbiome of the human gut^6^), or a specific type of ecosystem (e.g., the core microbiome of a particular soil type^7^). However, much broader^8^ (e.g., the core microbiome of ruminant guts^9^) and much narrower (e.g., the core microbiome of a particular host population^10^ or host genotype^11^) habitat classes can be used as well. Because core microbial taxa are commonly, if not always, associated with a specific habitat, they are thought to be important for identifying microbial persistence requirements within the habitat, predicting ecosystem function^12–15^ or, in the case of host-associated microbiomes, understanding symbiotic relationships that may be important to host performance.^16–19^ By contrast, it is generally assumed that microbial taxa occurring only sporadically in a given habitat are more likely to be transients or other community members that do not contribute substantially to host or ecosystem function.

While the concept of a core microbiome is both simple and appealing, how to best define the core microbiome of any given habitat remains unclear.^1,2^ In general, three different core criteria have been proposed^1^ based on 1) the occupancy (i.e., incidence rate, frequency of occurrence) of microbial taxa across samples, 2) the abundance of microbial taxa in individual samples or across a set of samples, or 3) a combination of occupancy and abundance criteria.^20,21^ Within these three strategies, however, there are a wide range of parameters that can influence interpretation. One obvious example is the threshold occupancy or abundance that a microbial taxon must achieve to be considered part of the core. In some cases, microbial taxa are included in the core if they are found in at least two samples from a given habitat.^1^ In other cases, much higher core thresholds have been used, such as requiring that a microbial taxon be present in half or even all of the samples from a particular habitat. Some studies even present analyses across a range of core thresholds so that results can be interpreted in context.^22,23^

Beyond the occupancy and abundance thresholds, another decision that can influence identification of core microbial taxa is the taxonomic rank or phylogenetic level at which the core microbiome is defined. Studies exist spanning a wide range of taxonomic ranks and phylogenetic levels from phyla to genera and from 97% operational taxonomic units (OTUs) to amplicon sequence variants (ASVs).^1^ Not surprisingly, examining the core microbiome at different taxonomic and/or phylogenetic scales often results in different conclusions. Higher taxonomic ranks like phyla and order tend to be more conserved across samples, resulting in a larger proportion of microbial taxa being identified as part of the core. However, ecology and function tend to be less conserved at these higher taxonomic ranks^24,25^, making the core microbiome less useful for informing host or ecosystem ecology. By contrast, at very low taxonomic ranks, it is unclear whether variation between microbial taxa has significant ecological or functional consequences. ASVs, for example, may only differ by one base pair in the portion of the marker gene used for sequencing.^26^ However, in microbial taxa with multiple copies of the 16S rRNA marker gene, individual gene copies can vary intragenomically by >1%.^27,28^ Thus, ASVs may be much more diverse than the populations or functions that they are being used to represent.^29^ This can impact the core microbiome in one of two ways. First, if multiple closely related ASVs are present in a threshold number of samples, they may all be counted individually as part of the core, inflating core diversity. Second, if individual samples each contain unique but closely related ASVs, none of the ASVs may occur in sufficient samples to be included in the core, reducing core diversity. Perhaps because of the challenges associated with defining core microbiomes at very high and very low taxonomic and/or phylogenetic scales, most analyses of core microbiomes have used intermediate scales, for example 97% operational taxonomic units (OTUs) or genera.^1^

The importance of taxonomic and/or phylogenetic scale, and the apparent variation in study outcomes as a function of the taxonomic/phylogenetic scale considered are not unique to analysis of core microbiomes.^1^ Indeed, these same challenges occur in more general types of ecological analyses of both microbial and macrobial systems, for example, calculation of ɑ- and β-diversity in and between ecological communities. This has led to the development of different phylogeny-aware metrics including Faith’s phylogenetic diversity^30^ (PD), Rao’s quadratic entropy^31^ and phylogenetic entropy^32^ for ɑ-diversity, and UniFrac distances^33^ for β-diversity. Phylogeny-aware metrics are advantageous because they integrate information across taxonomic ranks based on the branch lengths in a phylogenetic tree. This removes the need to select an arbitrary taxonomic rank or phylogenetic level for any particular ecological analysis. Further, because some phylogenetic properties can serve as indicators of both functional diversity^34^ and evolutionary potential^35^ (i.e., capacity to evolve in response to a perturbation, for example environmental change^36–38^), phylogeny-aware metrics often provide more informative comparisons^39^ of properties like diversity than do analyses at any single taxonomic or phylogenetic scale.

Despite the many benefits of phylogeny-aware ecological analyses, few phylogeny-aware methods for characterizing the core microbiome exist. Several recent papers have begun to account for phylogeny when characterizing core microbiomes^40–42^, for instance by defining core taxa based on a series of clustering thresholds.^43^ Nevertheless, there remains a lack metrics that smoothly integrate phylogenetic information across taxonomic and phylogenetic scales. In this paper, we develop a framework for analyzing core microbiomes using phylogenetic methods. To do this, we first develop the concept of a ‘core community phylogeny’ - a phylogeny where branches are defined based on being present in multiple samples from a single type of host or environment. We then use this phylogeny to examine common measures of core communities, including Venn diagrams, as well as metrics of within- and between-habitat diversity.

### Core Community Phylogenies

All phylogeny-aware metrics of ecological communities, for example Faith’s phylogenetic diversity (PD) and UniFrac distance, are based on an underlying phylogenetic tree comprising taxa present in the community. Logically, the analogous phylogenetic tree for core microbiomes should comprise all microbial taxa that are conserved across microbiomes from a particular type of habitat. However, for core microbiomes, there are two possible ways of defining such a tree: a ‘tip-based’ approach, and a ‘branch-based’ approach.

#### Tip-based phylogeny

One way to build a core community phylogeny is to first identify core microbial taxa based on their relative abundance and/or occupancy across a threshold number of microbiome samples from a particular type of habitat. Once these microbial taxa have been identified, a phylogenetic tree can be reconstructed based on the sequences (or other traits) of the conserved taxa. Notably, conserved microbial taxa can be identified at any pre-selected taxonomic rank (e.g., phyla, genera, 97% OTUs, ASVs, etc.) provided that a phylogenetic tree can be built connecting the various microbial lineages. The tip-based phylogeny is useful in that it incorporates phylogeny while still identifying extant microbial taxa that comprise the core. However, because the tip-based phylogeny identifies core taxa without reference to their phylogenetic relationships, it does not capitalize fully on phylogenetic information. Consequently, metrics calculated using the tip-based phylogeny are more susceptible to variation across taxonomic rank and/or phylogenetic level as compared to metrics calculated using the branch-based approach that we discuss next (see ‘Examples’ and ‘Discussion’ for more details).

#### Branch-based phylogeny

An alternative way to build a core community phylogeny is to first build a phylogeny of all microbial taxa present in a set of microbiome samples from a particular type of habitat. This phylogenetic tree can then be examined, branch-by-branch, to determine which branches satisfy core criteria based occupancy and/or abundance thresholds (i.e., which branches occur in sufficient numbers of samples or have sufficient abundance associated with them). Branches that satisfy the core criteria can then be retained, while branches that do not can be discarded. Note that, at least for a rooted tree, this results in discarding shallow branches connected to deeper branches that have also been discarded, such that the remaining phylogeny is fully connected (but see below for issues that arise with unrooted trees). Like the tip-based phylogeny, the branch-based phylogeny can have leaf nodes at any taxonomic rank. However, because the branch-based phylogeny integrates information on shared microbial lineages across all depths of the phylogeny both for defining the core and calculating core metrics, there is less reason to use high taxonomic levels (see ‘Examples’ below). Thus, we recommend ASVs as the default choice.

Importantly, the tip-based phylogeny and the branch-based phylogeny for a given system will not always be identical. Whereas the branch-based phylogeny will always have all of the branches present in the tip-based phylogeny, the tip-based phylogeny will often be missing some of the branches present in the branch-based phylogeny (see Figure 1). More specifically, the tip-based phylogeny will be missing branches whenever a set of microbiome samples share closely related, but not identical, taxa. In this case, none of the shallow branches will be considered core for either the tip-based or the branch-based phylogeny. However, the deeper branches from which all of the related tips are derived may be present on the threshold number of samples and thus may be retained in the branch-based phylogeny. By contrast, these branches will be discarded in the tip-based phylogeny, since definition of core is solely based on shared leaf nodes. This fundamental difference is what makes the branch-based approach less sensitive to variations in the precise taxonomic scale considered, at least when making relative comparisons between two or more core communities (see ‘Examples’ below).

**Figure 1.**
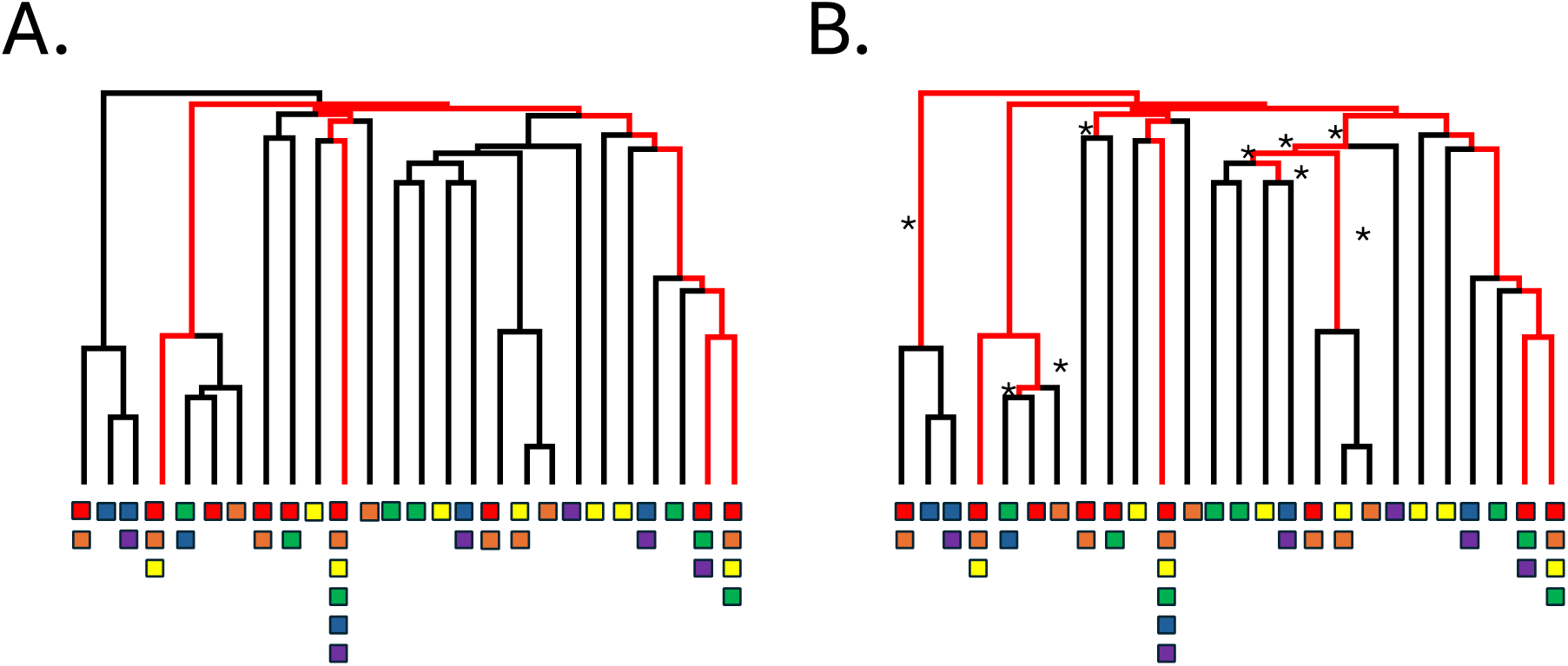
Examples of a tip-based phylogeny (A) and a branch-based phylogeny (B) for the same microbial phylogeny. Branches of the microbial phylogeny that are not part of the core community phylogeny are shown in black, while branches that are part of the core community phylogeny are shown in red. For this example dataset, there are 6 samples (red, orange, yellow, green, blue, and purple squares) and we use an occupancy threshold of 0.5 as our core criterion. Thus being part of the core requires being present in 3 samples. Colored boxes at the end of each terminal branch indicate the samples that a particular microbial taxon is present on. Stars are used to mark branches that are included in the branch-based phylogeny but not in the tip-based phylogeny.

Figure 1 shows the construction of both a tip-based phylogeny (A) and a branch-based phylogeny (B) from the same set of microbiome samples and based on an occurrence threshold of 0.5 (i.e., a taxon must occur in at least 50% of samples to be considered part of the core). In Figure 1, the non-core branches of the microbial phylogeny are shown in black, while the core branches are shown in red. Notice that there are differences between the tip-based phylogeny and the branch-based phylogeny (see starred branches). In particular, the branch-based phylogeny includes additional internal branches that are missing from the tip-based phylogeny. This occurs because some clades are represented in 50% of samples, and thus are part of the branch-based core microbiome, even though the individual taxa within the clades are not present in 50% of samples.

#### Rooted versus Unrooted Trees

An important consideration in defining the core community phylogeny is whether to assume that the root node is the most recent common ancestor (MRCA) of all taxa in the sample. The root node should only be used as the MRCA when a rooted tree is provided. Otherwise, the root node is arbitrary, and there is no reason to believe that the root node is any more basal than the other nodes in the tree. Fortunately, microbial phylogenies generated using pipelines like Qiime2^44^ usually have the option for rooting trees (although this is typically done through midpoint rooting, which involves multiple assumptions). Similar to UniFrac, we strongly encourage using a rooted tree and forcing the root node to be the MRCA. This is particularly true for branch-based core community phylogenies. The reason is as follows: using the branch-based approach, branches are counted based on their inclusion in individual phylogenies from each sample. When a root node is defined, these individual sample phylogenies necessarily include the root. In absence of a root, however, individual sample phylogenies only span the taxa present in each sample (core or otherwise). Consequently, basal branches may not be counted if entire clades are differentially distributed across subsets of samples. When this happens, distinct clades are not necessarily connected in the phylogeny that emerges. To remedy this, we connect the final core community phylogeny based on the MRCAs for all of the distinct clades. This, however, does not guarantee that branches will not be lost in the process. For tip-based phylogenies, the choice to use an unrooted tree is less problematic because the branches of the core community phylogeny are selected based on the occupancy of microbial taxa (tip nodes), not on the occupancy of the branches themselves. Even for tip-based trees, however, we recommend using a rooted tree and assuming that the root node is the MRCA.

#### Criteria for Inclusion

Another important consideration in defining the core community phylogeny is the criterion used to determine whether a taxon or branch is part of the core. The simplest option is to use an occupancy threshold. Core taxa could, for example be selected based on being present in 50%, 90% or 100% of samples from a particular type of habitat. Alternatively, various relative abundance thresholds could be used, either by themselves or in combination with an occupancy threshold. Core taxa could, for instance, be selected based on attaining a mean or minimum relative abundance across all samples or based on reaching a maximum relative abundance in at least one sample. Whereas occupancy thresholds imply a degree of conservation across samples, abundance thresholds indicate numerical dominance (with the caveat that measured dominance may be different from true dominance due to biases in extraction, PCR, etc.). When used in combination, occupancy and abundance thresholds allow the core to be selected based on both conservation and numerical dominance.

While occupancy and abundance thresholds are the simplest forms of core criteria, several other definitions have been developed as well. Recently, for instance, Shade and Stopnisek^20^ have pioneered a method that identifies core taxa based on being both strongly conserved within a habitat and contributing to variation across that habitat. Fundamentally different from the threshold approach, this more nuanced method can, in some cases, be more powerful than simple occupancy and abundance thresholds. Importantly, identification of a core community phylogeny does not depend on the use of any particular core criterion. Rather, core community phylogenies can, at least theoretically, be defined based on any criterion that can be applied to individual taxonomic units. For the tip-based tree, this means applying the chosen criterion when selecting core taxa. For the branch-based tree, this means applying the chosen criterion on a branch-by-branch basis.

### Phylogenetic Metrics for Core Microbiomes

Once a core community phylogeny has been constructed, it can be used to perform phylogenetic analyses of microbiome samples from different types of habitats. Here we consider several common analysis methods, including Venn diagrams, a core microbiome equivalent of Faith’s phylogenetic diversity (α-diversity), and a core microbiome equivalent of UniFrac distance (*β*-diversity).

#### Venn Diagrams

Venn diagrams are one of the most common methods for visualizing core microbiomes across habitats. Non-phylogenetic Venn diagrams show counts of the core microbial taxa shared by different combinations of habitat types. This can be extended to a phylogenetic framework by replacing counts of shared microbial taxa with shared phylogenetic branch lengths. Briefly, the core community phylogenies from different habitat types can be compared, branch-by-branch, to determine which branches appear in which habitat combinations. The total lengths of these shared branch segments can then be calculated and reported either as an absolute value or as percentages of the total branch length spanning the core community phylogenies of all habitats in the analysis.

#### Core Faith’s Phylogenetic Diversity (PD)

The phylogenetic equivalent to richness (a count of microbial taxa present in a community), Faith’s PD is the sum of the branch lengths of all branches spanning the taxa present in a particular microbiome sample or ecological community. This definition can easily be extended to core microbiomes by summing the branch lengths of all branches in the core community phylogeny. Doing so gives a measure of how much phylogenetic diversity is conserved across microbiome samples from a particular type of habitat.

#### Core UniFrac Distances

The phylogenetic equivalent to Jaccard’s index^45^ (the fraction of unique microbial taxa between two communities), the UniFrac distance^33,46^ between two communities is the sum of the branch lengths that are unique to each microbiome sample or ecological community divided by the sum of the branch lengths of all branches spanning taxa present in both microbiome samples or ecological communities. Again, this definition can be extended to core microbiomes by dividing the branch lengths unique to each core community phylogeny by the total branch lengths of the two core community phylogenies combined. Doing so gives a measure of how phylogenetically distinct the core microbiomes are from two different types of habitats.

### Examples

Additional file 1: Simple Example System (see Figures S1.1 and S1.2) outlines the calculation of Venn diagrams, core Faith’s PD, and core UniFrac distances using a set of highly simplified core community phylogenies. In this additional file, we show differences between tip- and branch-based approaches, and outline calculations with both rooted and unrooted trees. Throughout the remainder of the main text, we apply our methods to more realistic systems using rooted trees. First, we consider the *Staphylococcus* component of the human skin microbiota based on a dataset from Joglekar et al.^47^ For this analysis, we use the branch-based phylogeny combined with a simple occupancy threshold of 50% (i.e., a taxon/branch must be present in 50% of samples to be considered part of the core, for alternate core criteria see Additional file 1: Additional *Staphylococcus* Analyses, Figures S2.1-S2.3). Figure 2 shows *Staphylococcus* taxa present in skin samples from two different people, first based on amplicon sequence variants (ASVs) and then based on operational taxonomic units (OTUs, see ‘Methods’ for more details). Core taxa that would be identified based on a non-phylogenetic approach are identified with black stars. Core branches identified based on our phylogenetic approach are colored red. The *Staphylococcus* microbiota on these two individuals demonstrate the challenges faced by non-phylogenetic definitions of the core microbiome: whereas Individual One (Fig. 2A) has a lower core *Staphylococcus* diversity based on ASVs (3 versus 6), Individual Two (Fig. 2B) has a lower core *Staphylococcus* diversity based on OTUs (3 versus 4). Two separate phenomena explain this diversity reversal. First, on Individual One, there are two ASVs from the CCE Clade (see Figure 2A) that do not, independently, reach an occupancy threshold of 50% and thus fail to contribute to core ASV richness. When these ASVs are pooled into a single OTU, however, the OTU is present in 50% of samples and thus is counted as part of the core. This leads to an increase in diversity at the OTU scale and reflects the fact that strain variation across samples from Individual One obscured the relatively high occupancy of closely related *Staphylococcus* in the CCE clade. Second, on Individual Two, three closely related *S. capitis/caprae* strains and two closely related *S. hominis* strains separately contribute to core ASV richness. These ASVs are, however, pooled and counted as single core taxa at the OTU scale. This leads to a reduction in OTU diversity and reflects the fact that most of the ASV diversity on Individual Two was a result of shallow strain variation within the *S. capitis/caprae* and *S. hominis* clades. Unlike non-phylogenetic measures of core *Staphylococcus* diversity, phylogenetic measures do not vary as much across phylogenetic levels. Rather, phylogenetic measures suggest that the core microbiota of the first person is more diverse at both the ASV (0.169 vs. 0.113) and OTU (0.180 vs. 0.105) scales. The explanation is simple: the relatively fewer core ASVs on Individual One span a wider phylogenetic breadth than the numerous but closely related ASVs on Individual Two (See Additional file 1: Additional *Staphylococcus* Analyses for further comparisons across individuals from the Joglekar et al.^47^ study). Rank reversals in diversity estimates as a function of phylogenetic scale can be problematic because they can make it difficult to interpret differences between treatment groups. Using a non-phylogenetic approach, for instance, it might be difficult to assess whether or not core *Staphylococcus* diversity increases resistance to pathogens like methicillin-resistant *S. aureus* (MRSA).^48,49^ This is because the people considered to have high versus low core *Staphylococcus* diversity changes depending on the phylogenetic scale considered. By contrast, the stability of our phylogenetic metrics across all phylogenetic scales makes them more amenable to exploring relationships between core *Staphylococcus* diversity and disease resistance.

**Figure 2.**
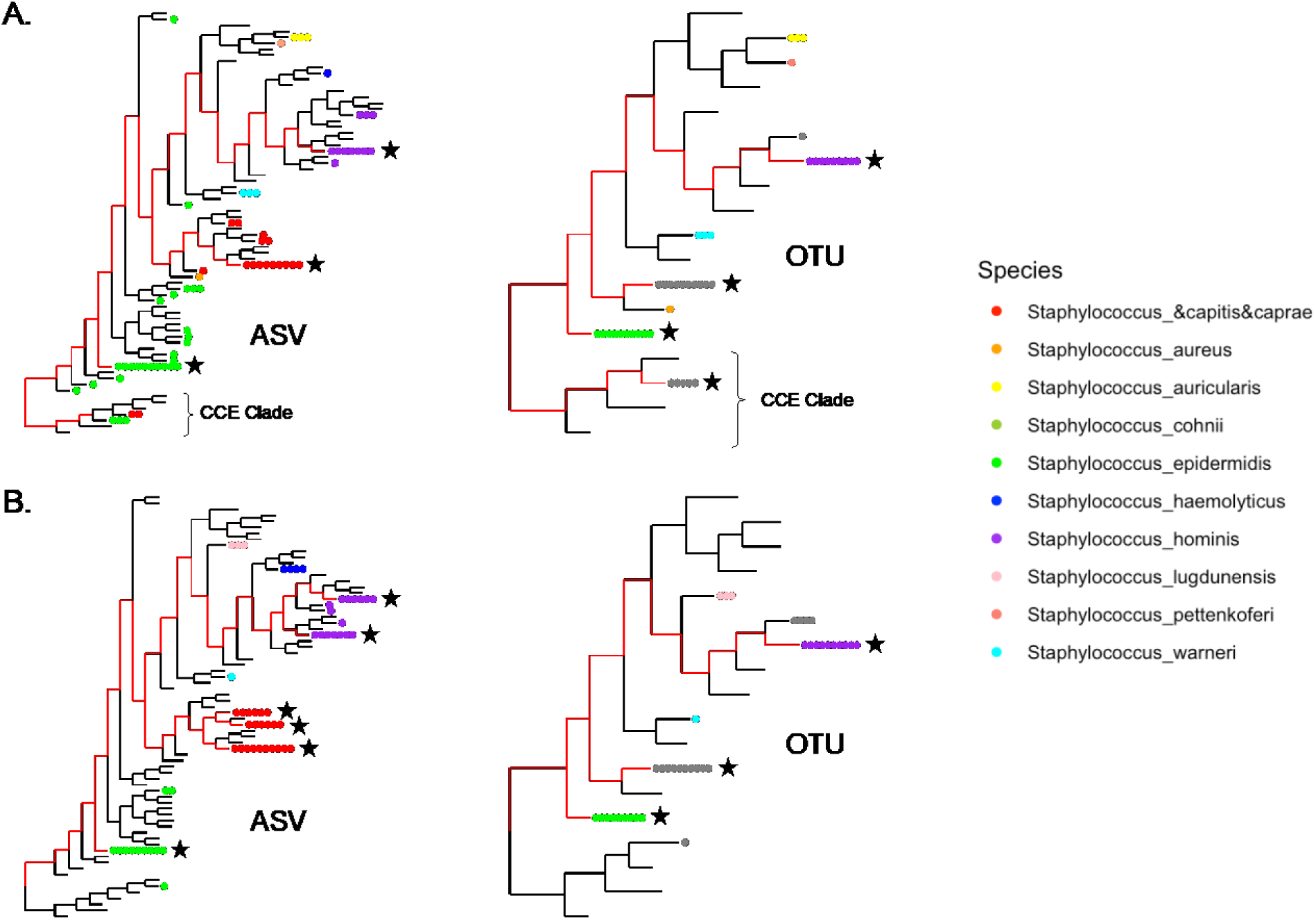
*Staphylococcus* phylogenies for 10 skin microbiome samples from two individuals - Individual One (A) and Individual Two (B) - using ASVs (left) and OTUs (right). Circles at the end of each terminal branch indicate the number of samples that contained the corresponding ASV or OTU. These circles are colored based on assigned species, with grey being used when multiple species are contained within the same OTU. Black stars indicate taxa that would be part of each individual’s core microbiome based on a non-phylogenetic approach using an occupancy threshold of 50%. Red lines indicate branches that are part of each individual’s core microbiome based on a branch-based phylogenetic approach using the same occupancy threshold of 50%. Because the terminal branches on these trees are extremely short, trees are visualized using a square root transformation. This transformation was not used in calculating core microbiota.

Figure 3 shows a comparison of the core *Staphylococcus* microbiota across skin habitats from the same Jonglekar et al.^47^ study, along with turnover (β-diversity) in core *Staphylococcus* composition among habitats. Again, differences in the phylogenetic and non-phylogenetic approaches are apparent. Based on the non-phylogenetic Jaccard distance, for instance, the core *Staphylococcus* microbiota of the nares is highly distinct from the core *Staphylococcus* microbiota of all other skin habitats. This distinction, however, is much less obvious when using a tip-based UniFrac distance and non-existent when using a branch-based UniFrac distance. Indeed, the branch-based UniFrac distance suggests that the core *Staphylococcus* microbiota of the nares is actually quite similar to the core *Staphylococcus* microbiota of both moist and sebaceous sites. Inspection of the phylogenetic trees for the tip- and branch-based approaches explains these differences. In particular, the nares are missing an *S. hominis* ASV found in the core *Staphylococcus* microbiota all other skin habitats. The nares are not, however, entirely lacking *S. hominis* ASVs. Rather, the nares are colonized by a range of other *S. hominis* ASVs, though none individually reaches the threshold for core inclusion. Nevertheless, because multiple *S. hominis* ASVs are present in nares samples, the branch-based approach counts most of the diversity associated with the *S. hominis* clade towards the core microbiota of both the nares and the other four habitat classes (sebaceous, moist, dry, feet). It is only the shallow tip branches that do not contribute towards the core microbiota of the nares. By contrast, for the tip-based and non-phylogenetic approaches, the missing *S. hominis* ASV drives significant turnover between the nares and all other sites.

**Figure 3.**
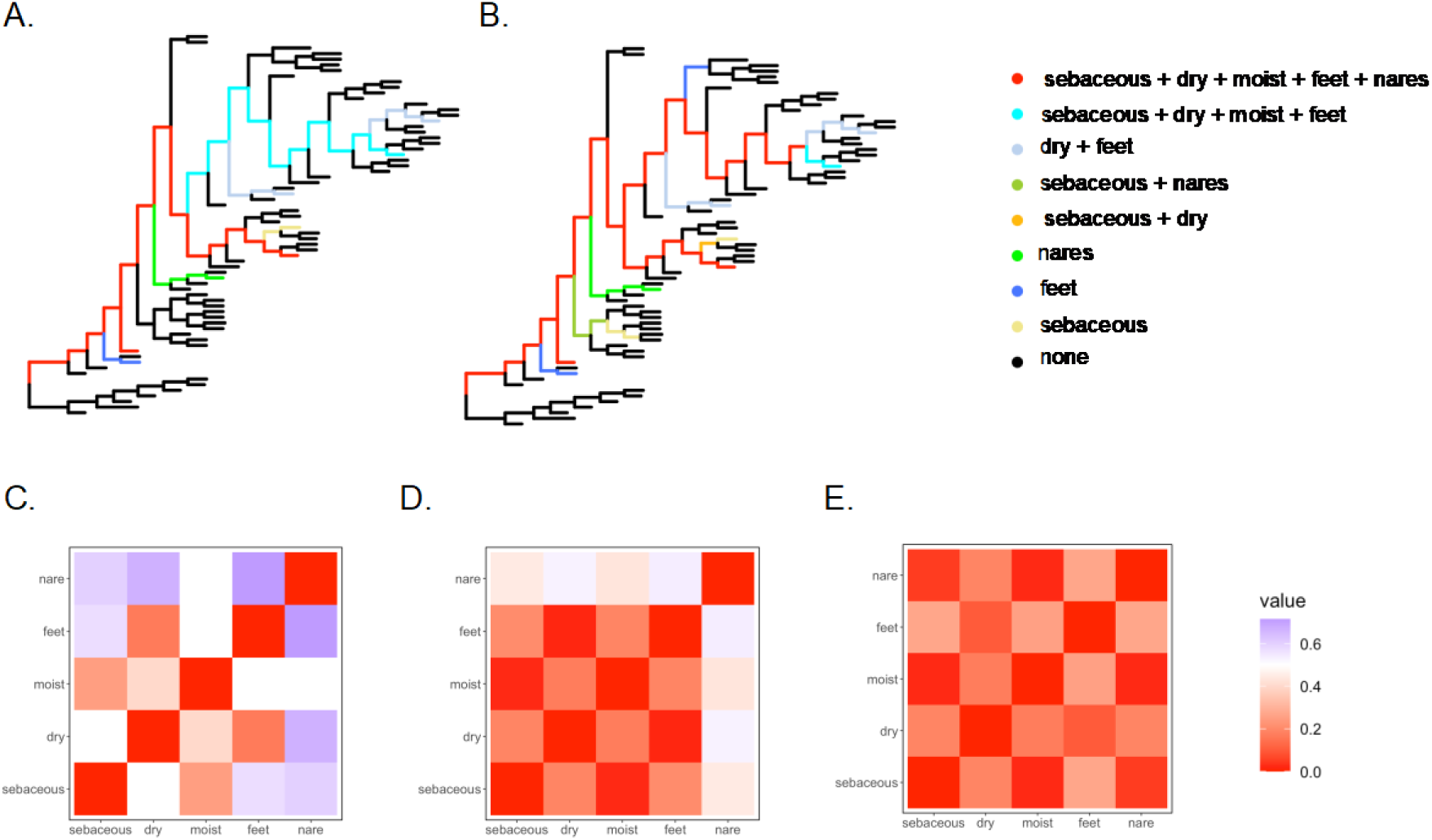
(A) Tip-based and (B) branch-based *Staphylococcus* phylogenies showing the shared and unique core community phylogenies, along with Jaccard distances (C), tip-based UniFrac distances (D) and branch-based UniFrac distances (E) among skin habitats. All panels in this figure used a 50% occupancy threshold as the core criterion. Because the terminal branches on these trees are extremely short, trees are visualized using a square root transformation. This transformation was not used in calculating core microbiota.

Having demonstrated the benefits of our phylogenetic approach on a relatively simple and low diversity *Staphylococcus* system, we next turn our attention to two additional datasets – one host-associated and the other environmental – both of which contain significant diversity and multiple bacterial phyla. Specifically, we consider the gut microbiota of three different whiptail lizard (*Aspidoscelis*) species and the soil microbiota of three different soil types (see Materials and Methods). Figure 4 shows standard Venn diagrams based on ASV counts (A, D) along with our new phylogenetic Venn diagrams for tip-based (B, E) and branch-based (C, F) trees. For all Venn diagrams, we define the core based on a 50% occupancy threshold (for alternate core criteria, see Additional file 1: Additional Venn Diagrams). As compared to our phylogenetic Venn diagrams, non-phylogenetic Venn diagrams (A, D) show the lowest percentage of system diversity in the compartments representing the intersections of (A) gut microbiota of all three lizard species (0%) and (D) microbiota of all three soil types (8.02%). Non-phylogenetic Venn diagrams also show the highest percentage of system diversity in (A) the three compartments unique to single lizard species (80.00%) and (D) the three compartments unique to individual soil types (76.88%). This is not surprising. Using a non-phylogenetic approach, microbial taxa from different host species/environments must be identical to be counted in intersection compartments. This biases against finding shared microbial diversity, particularly when using ASVs, because microbial lineages with small differences in 16S rRNA gene sequences (e.g., one base pair substitutions) are deemed unique. By contrast, with a phylogenetic approach, even if different host populations/environments have slightly different ASVs, this only contributes a small amount of diversity to the unique compartments of the Venn diagram. Meanwhile, the majority of the diversity is allotted to intersection compartments because the ASVs from the different host populations/environments share most of their phylogenetic history. Comparing branch-based and tip-based phylogenetic approaches, branch-based phylogenies show a higher percentage of system diversity in the compartments representing intersection across (B, C) all three lizard species (27.10% vs. 19.16%) and (E, F) soil types (29.41% vs. 27.14%). This reflects the inclusion of additional, deep internal branches in branch-based phylogenies that are dropped from tip-based phylogenies (see Figure 1). These basal branches are more likely to be conserved across different host species/environments (i.e., phyla are more likely to be shared than ASVs), inflating diversity in shared compartments of the Venn diagram (see Additional file 1: Additional Venn Diagrams Figures S3.1-S3.4 for several additional datasets, including lizard skin and lake water).

**Figure 4.**
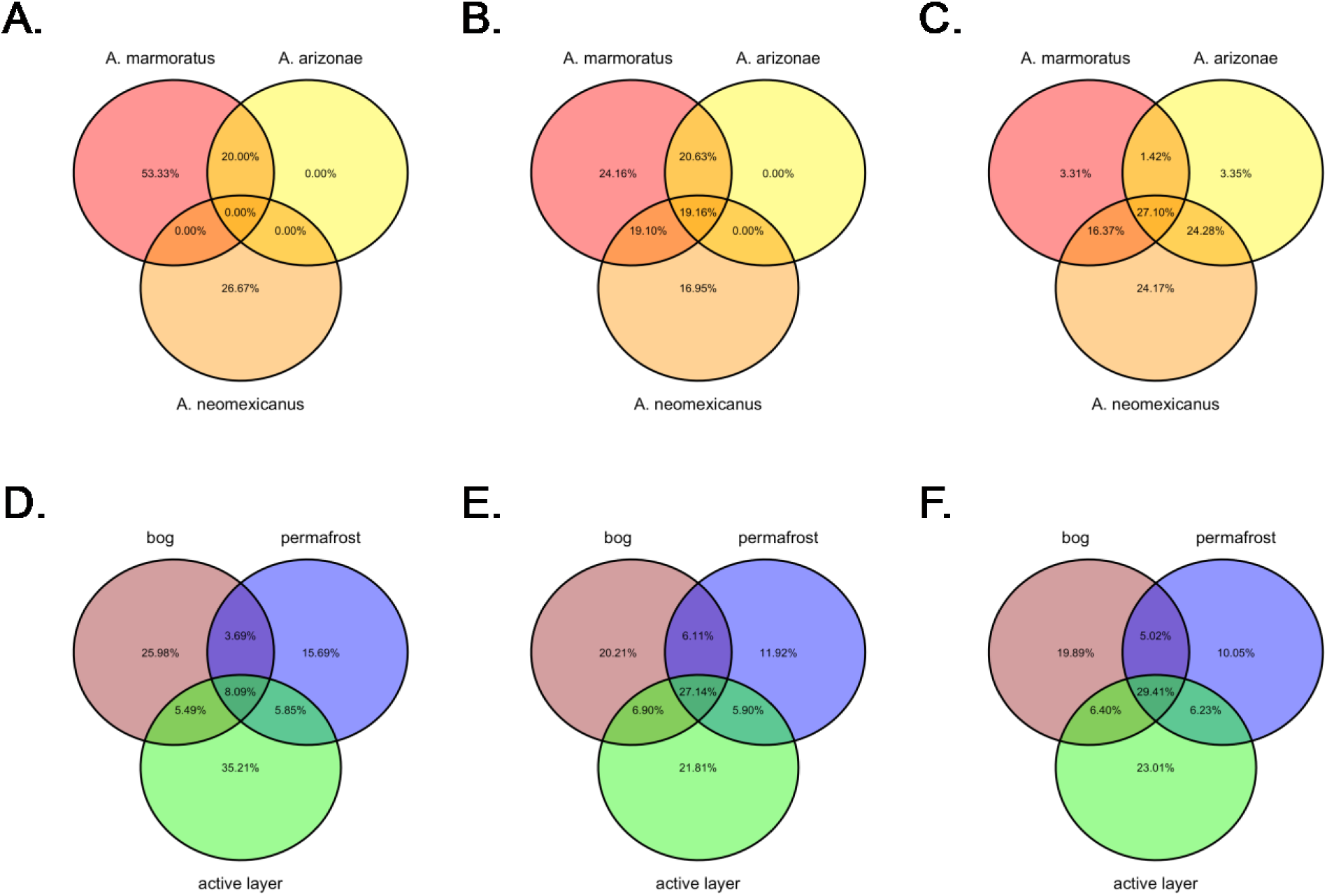
Venn diagrams for lizard gut (A-C) and soil (D-F) microbiota showing the percentage of the total ASV counts shared by the core microbiota from each lizard species or soil type (A, D) and the phylogenetic branch length shared by the core microbiota from each lizard species or soil type using either tip-based (B, E) or branch-based (C, F) phylogenies. In all Venn diagrams, lizard populations and soil types colored are as follows: *Aspidoscelis arizonae* (yellow), *A. neomexicanus* (orange), *A. marmoratus* (red), bog soil (brown), permafrost soil (blue) and seasonally thawed active layer soil (green).

Figure 5 (A-C) shows core richness, and tip- and branch-based core Faith’s PD as a function of taxonomic rank for lizard gut microbiota based on an occupancy threshold (A), an occupancy and abundance threshold (B) and the Shade and Stopnisek core criterion (C). For the sake of comparison across metrics and taxonomic rank, we have plotted core diversity of two lizard species (yellow: *A. arizonae*; red: *A. marmoratus*) relative to the third lizard species (*A. neomexicanus*; see Materials and Methods). Figure 5 (E-G) shows similar plots for soil microbiota, this time agglomerating taxa based on cophenetic distance, rather than taxonomy. Again, for the sake of comparison, we have plotted core diversity of bog (brown) and active layer (green) soils relative to permafrost soils. Figure 5 (D, H) shows the range of values obtained for core richness and tip- and branch-based Faith’s PD across all taxonomic ranks/tip agglomerations considered. In most cases, phylogenetic measures of lizard gut and soil microbiota show less variation across taxonomic rank/tip agglomeration as compared to non-phylogenetic measures. Using an occupancy threshold of 50%, for instance, the richness of the core gut microbiota of *A. marmoratus* (red) is anywhere from 0.64 (microbial species) to 2.75 (microbial ASV) times as diverse as the core gut microbiota of *A. neomexicanus*. By contrast, for the same occupancy threshold, the branch-based core Faith’s PD suggests a much narrower range from 0.52 (microbial ASV) to 0.62 (microbial order). Likewise, using the Shade and Stopnisek criterion, the richness of the core microbiota of active layer soil is anywhere from 0.92 (agglomeration: h = 0.15) to 1.5 (ASV) times as diverse as the core microbiota of permafrost soil. Meanwhile, for the same Shade and Stopnisek criterion, the branch-based core Faith’s PD exhibit a much narrower range from 1.19 (agglomeration: h = 0.15) to 1.22 (ASV). Not all differences in metric stability across taxonomic rank/tip agglomeration are as dramatic. Still, the branch-based core Faith’s PD typically shows the least variation across taxonomic ranks/tip agglomeration, while the tip-based core Faith’s PD shows the second least variation, and richness shows the most variation. Nevertheless, there are deviations to this general trend, particularly for the Shade and Stopnisek criterion (see Additional File 1: Additional α- and β-diversity Analyses Figures S4.1 and S4.2 for several additional datasets, including lizard skin and lake water).

**Figure 5.**
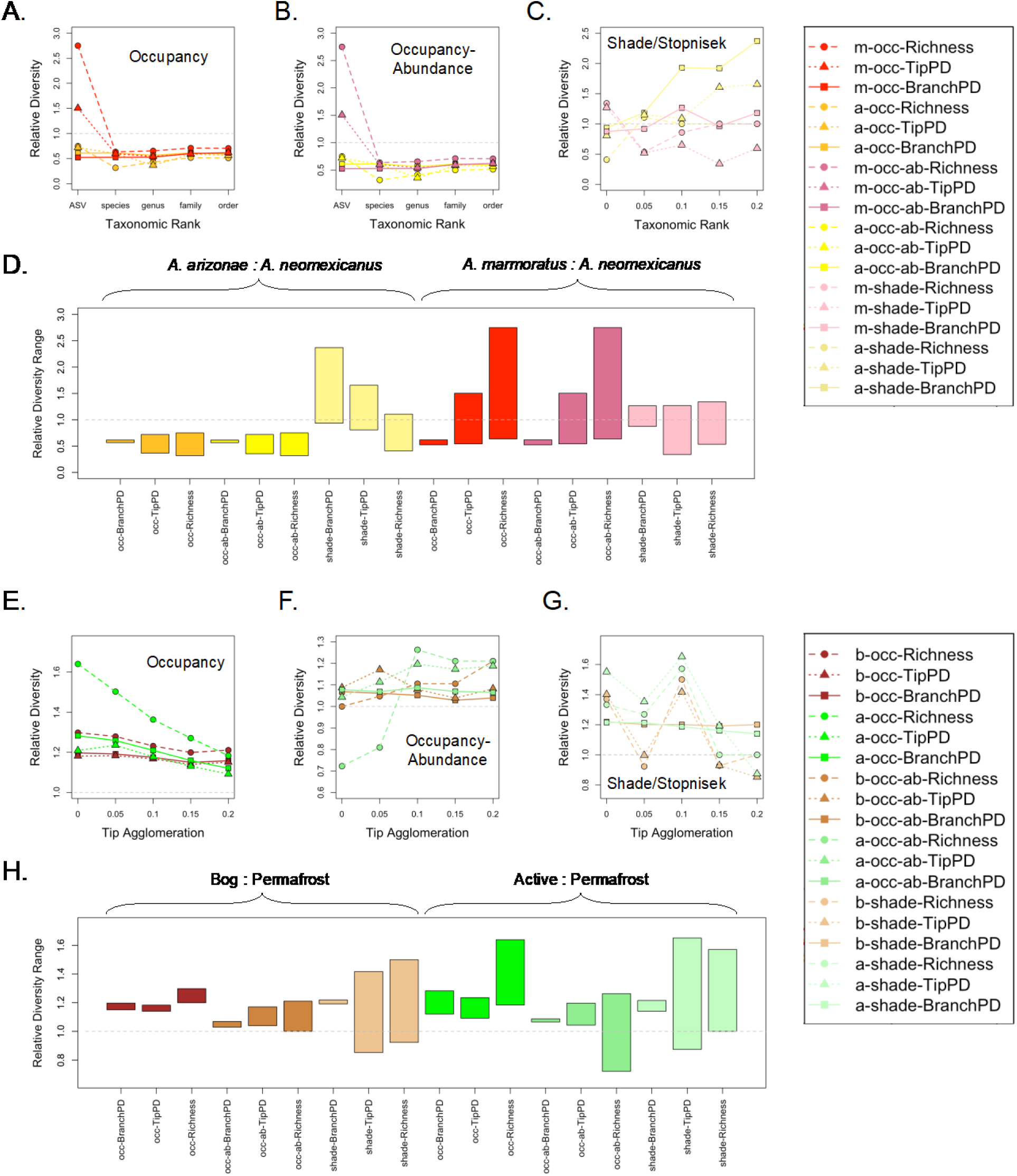
Comparison of alpha diversity across taxonomic ranks (A-C) and degrees of tip agglomeration (E-G, see Materials and Methods) for the branch-based Core Faith’s PD (solid, squares), the tip-based Core Faith’s PD (dotted, triangles), and richness (dashed, circles) assuming the following core criteria: (A,E) an occupancy threshold of 50% (gold, red, dark brown, bright green), (B,F) an occupancy threshold of 50% and a mean abundance threshold of 0.1% (yellow, dark pink, medium brown, medium green), (C,G) the Shade and Stopnisek method (light yellow, light pink, light brown, light green). In (A-C), we show the core diversity of *Aspidoscelis arizonae* (gold, yellow, pale yellow) and *A. marmoratus* (red,dark pink, light pink) gut microbiota relative to the core diversity of *A. neomexicanus* gut microbiota. In (E-G) we show the core diversity of bog (dark, intermediate and light brown) and active layer (bright, intermediate and light green) soil microbiota relative to the core diversity of permafrost soil microbiota. Bar graphs (D,H) showing minimum and maximum values of alpha diversity across taxonomic rank for *A. arizonae* (gold, yellow, light yellow) and *A. marmoratus* (red, dark pink, light pink) relative to *A. neomexicanus*, and across tip agglomeration for bog soil (dark, intermediate and light brown) and active layer soil (bright, intermediate and light green) relative to permafrost soil. Diversity metrics are indicated as follows: ‘BranchPD’ is the branch-based phylogeny approach, ‘TipPD’ is the tip-based phylogeny approach, and ‘Richness’ is a count of taxa for each taxonomic rank. Core criteria are indicated as follows: ‘occ’ is the 50% occupancy threshold, ‘occ-ab’ is the 50% occupancy and 0.1% abundance threshold, and ‘shade’ is the Shade and Stopnisek method.

Figure 6 (A-C) shows the Jaccard, tip- and branch-based UniFrac distances between core gut microbiota of *A. neomexicanus* and both *A. arizonae* (yellow) and *A. marmoratus* (red) based on an occupancy threshold (A), an occupancy and abundance threshold (B) and the Shade and Stopnisek core criteria (C). Figure 6 (E-G) shows similar plots for soil microbiota, this time agglomerating taxa based on cophenetic distance, rather than taxonomy. Figure 6 (D, E) shows the range of values obtained for Jaccard, tip- and branch-based UniFrac distances across all taxonomic ranks/tip agglomerations considered. Broadly speaking, the branch-based approach gives the smallest distances, as well as the least variation across taxonomic rank/tip agglomeration. Meanwhile the tip-based approach gives intermediate distances and intermediate variation and the non-phylogenetic approach (i.e., Jaccard distance) gives the largest distances and the most variation (see Additional file 1: Additional α- and β-diversity Analyses Figures S4.1 and S4.2 for several additional datasets, including lizard skin and lake water).

**Figure 6.**
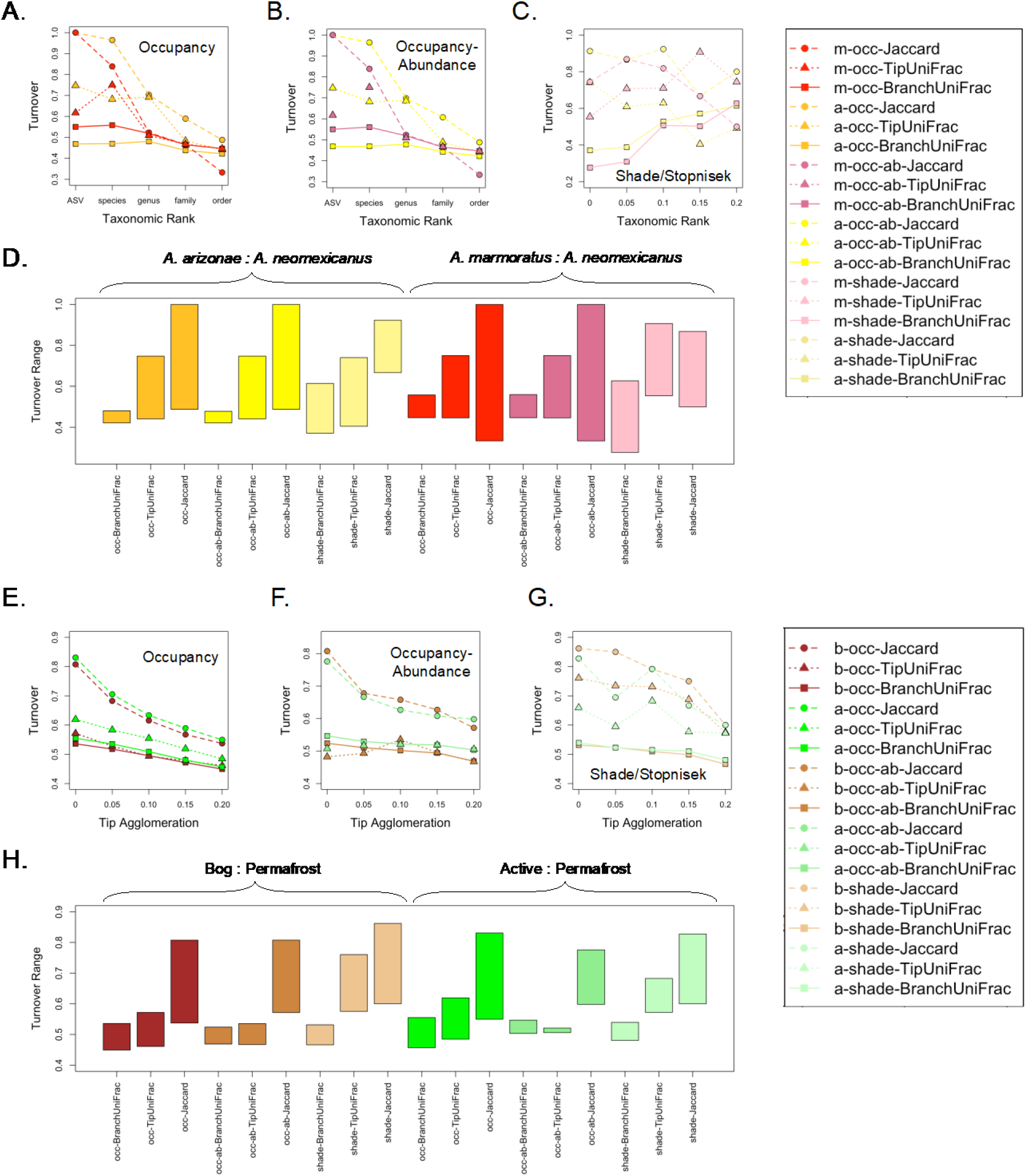
Comparison of beta diversity across taxonomic ranks (A-C) and degrees of tip agglomeration (E-G, see Materials and Methods) for the branch-based Core UniFrac distance (solid, squares), the tip-based Core UniFrac distance (dotted, triangles), and Jaccard distance (dashed, circles) assuming the following core criteria: (A,E) an occupancy threshold of 50% (gold, red, dark brown, bright green), (B,F) an occupancy threshold of 50% and a mean abundance threshold of 0.1% (yellow, dark pink, medium brown, medium green), (C,G) the Shade and Stopnisek method (light yellow, light pink, light brown, light green). In panels (A-C), we show turnover of *A. arizonae* (gold, yellow, light yellow) and *A. marmoratus* (red, dark pink, light pink) gut microbiota relative to *A. neomexicanus* gut microbiota. In panels (E-G) we show turnover of bog (dark, intermediate and light brown) and active layer (bright, intermediate and light green) soil microbiota relative to permafrost soil microbiota. Bar graphs (D,H) showing minimum and maximum values of beta diversity (turnover or distance) across taxonomic rank for *A. arizonae* (gold, yellow, light yellow) and *A. marmoratus* (red, dark pink and light pink) relative to *A. neomexicanus* and across tip agglomeration for bog soil (dark, intermediate and light brown) and active layer soil (bright, intermediate and light green) relative to permafrost soil. Diversity metrics are as follows: where ‘BranchUniFrac’ is the branch-based phylogeny approach, ‘TipUniFrac’ is the tip-based phylogeny approach, and ‘Jaccard’ is a count of taxa for each taxonomic rank/tip agglomeration. Core criteria are indicated as follows: ‘occ’ is the 50% occupancy threshold, ‘occ-ab’ is the 50% occupancy and 0.1% abundance threshold, and ‘shade’ is the Shade and Stopnisek method.

### The holobiont R package

To facilitate the use of our new phylogenetic measures of core microbiomes, we developed a new R package, `holobiont`, that includes functions for identifying the core community phylogeny in any microbiome, drawing phylogenetic Venn diagrams, calculating the core Faith’s PD for a set of communities, and calculating the core UniFrac distance between two sets of communities. Briefly, the `coreEdges` function takes a phyloseq object^50^ and the core criterion and identifies the branches of the core community phylogeny. The `coreTree` function takes the same information and colors the branches of the core community phylogeny red, while leaving the remaining branches of the community tree black. The `corePhyloVenn function` takes a phyloseq object, a vector defining the habitat class for each microbiome sample, and the core criterion and draws a phylogenetic Venn diagram of either branch lengths or percentages of branch lengths shared by the core community phylogenies of different combinations of habitat classes. The `coreVennTree function` takes the same input as the `corePhyloVenn` function and colors the branches of the phylogenetic tree based on the combination of habitat class core community phylogenies that they belong to. The `coreFaithsPD` function takes a phyloseq object and the core criterion and calculates the core Faith’s PD. Finally, the `coreUniFrac` function takes a phyloseq object and a vector specifying which samples belong to which habitat class (maximum 2) and calculates the UniFrac distance between the core community phylogenies. In addition to the phylogenetic functions, the `holobiont` package also provides corresponding non-phylogenetic functions, including `coreTaxa, coreVenn, coreRichness` and `coreJaccard.`

Additional details outlining the methods and use of our R package can be found on the CRAN website at https://cran.r-project.org/web/packages/holobiont/index.html. A brief discussion of our modified Shade and Stopnisek algorithm can be found in Additional file 1: Shade and Stopnisek Algorithm.

## Discussion

In this paper, we propose a framework for performing phylogeny-aware analyses of core microbiomes. More specifically, we develop the concept of a core community phylogeny - a phylogeny based on the microbial lineages that are core to a specific habitat class. We then propose two different methods for selecting this tree - a tip-based approach and a branch-based approach. For the tip-based approach, core membership is decided based on occupancy and/or abundance of the leaf (tip) nodes. This is akin to the standard, non-phylogenetic method for defining a core microbiome. However, once members of the core microbiome have been identified, a phylogenetic tree is constructed. This allows for calculation of phylogeny-aware metrics that leverage the relatedness of the leaf nodes to determine overall properties (e.g., diversity) of the core microbiome. For the branch-based approach, core membership is decided independently for each branch of the phylogenetic tree. Therefore, in the branch-based approach, phylogeny informs not only the calculation of metrics but also, the identification of the core microbiome itself. The downside of the branch-based approach, however, is that the ‘core microbiome’ often includes internal nodes while excluding their associated leaf nodes. Thus, metrics calculated using the branch-based approach are not necessarily based on a set of extant taxa. Rather, they are based on a set of complete and partial microbial lineages (see, for example, Figure 2A, where branches leading to the CCE clade are included in the branch-based core phylogeny, even though all terminal branches from this clade are excluded). Still, because the branch-based approach uses phylogenetic information to both define the core microbiome and to calculate core metrics, we suggest using the branch-based approach except when there is a strong desire to identify specific microbial taxa within the core (e.g., to select for culturing experiments).

As compared to existing metrics, the primary advantage of our phylogenetic metrics is that they seamlessly integrate core microbiome properties (i.e., richness, turnover, shared taxa) across taxonomic scales/phylogenetic levels. Thus, our phylogenetic metrics are more likely to provide biologically relevant information. This is especially true when non-phylogenetic metrics are calculated at taxonomic ranks that fail to capture meaningful biological variation.^51^ If, for example, the chosen taxonomic rank is too low, substantial biological variation may be identified that has no functional or ecological consequences. Alternatively, if the chosen taxonomic rank is too high, large amounts of meaningful biological variation may be pooled into the same taxonomic unit. Depending on the system, the type of variation considered and the metric calculated, it can be difficult to *a priori* determine (or even *a posteriori* assess) which taxonomic ranks are more or less likely to be biologically informative. Further, in certain instances multiple taxonomic ranks exhibit substantial meaningful biological variation. In this case, a non-phylogenetic metric at any single taxonomic rank - even a well-chosen taxonomic rank - cannot fully capture the underlying signal. Phylogenetic metrics circumvent these challenges by summarizing the full spectrum of biological variation across all taxonomic ranks, with variation at lower taxonomic ranks weighted less than variation at higher taxonomic ranks.

Another benefit of our branch-based approach (but not our tip-based approach) is that it overcomes challenges associated with defining the core microbiome in the face of high levels of strain variation. This is common in certain microbiome systems, including many host-associated microbiomes. The pervasive human skin isolate, *Staphylococcus epidermidis*, for example, evolves on individual humans over time,^52,53^ resulting in strain level variation across the human population. Another common human skin isolate, *Cutibacterium acnes*, exhibits even more dramatic strain-level variation, showing differences in sequences across pores from a single human individual.^54^ Indeed, in-depth profiling of the human microbiome has shown that there are numerous personalized microbial taxa that are found on only one or several individuals.^53,55^ Depending on the taxonomic rank at which the core microbiome is defined and the degree to which any single microbial strain is shared among individual hosts, natural strain-level variation can cause broadly shared microbial clades to be obscured from the non-phylogenetic core microbiome. By contrast, if we apply our branch-based approach to these same scenarios, only the terminal branches of the tree are omitted. Meanwhile, the more basal branches of the clades are retained because they are shared across all strains. We see this in our simple *Staphylococcus* example in Figure 2A, where the basal branches of the CCE clade are retained in the branch-based approach despite none of the extant ASVs from this clade reaching the non-phylogenetic threshold for core inclusion.

Because our phylogenetic metrics summarize microbiome measures across taxonomic rank/phylogenetic level and because the definition of the core microbiome - at least when using the branch-based approach - is less sensitive to strain-level variation, our phylogenetic metrics tend to be more consistent than comparable non-phylogenetic metrics when calculated at different taxonomic and/or phylogenetic scales. This is true both in a relative (i.e., comparison between microbiome samples) and an absolute sense. To see why phylogenetic metrics are more consistent, consider a habitat with hundreds of closely related core microbial strains within a single phylum. Using a non-phylogenetic approach, this system will have a very high ASV ɑ- diversity (i.e., ASV richness), but a very low phylum ɑ-diversity (i.e., phylum richness). By contrast, using our phylogenetic approach, ɑ-diversity (i.e., Faith’s PD) will be low regardless of whether the leaf nodes are defined at the ASV- or phylum-level. This is because, even when the leaf nodes are ASVs, all of the different ASV strains contribute very short branch lengths. Consequently, they do not substantially inflate ɑ-diversity beyond the diversity of the single phylum. The greater consistency of our phylogenetic metrics across taxonomic ranks means that it is less important to choose the correct taxonomic rank, and that it is rarely disadvantageous to use ASVs. This addresses one of the challenges that has been raised about interpreting core microbiomes when outcomes are variable across taxonomic rank or phylogenetic level.^1^

While our phylogenetic metrics typically yield lower variation across taxonomic rank and/or phylogenetic level, this is not always the case, particularly when using the Shade and Stopnisek criterion. Exactly why phylogenetic metrics are more variable when combined with the Shade and Stopnisek criterion is unclear. This may have to do with the emphasis that the Shade and Stopnisek method places on microbiome variation rather than microbiome conservation. Alternatively, it may be related to the stopping criterion used to delineate the Shade and Stopnisek core. Notably, changing the stopping criterion can impact the relative variation of phylogenetic and non-phylogenetic measures as a function of taxonomic rank (see Additional File 1: Shade and Stopnisek Algorithm Figure S5.1). While our phylogenetic metrics may not provide as much stabilization for the Shade and Stopnisek method, they still appear to perform as well or better in most cases. In our calculations of core α- and β-diversity for lizard guts and soils, for instance, our branch-based phylogenetic metric was more stable in 7/8 scenarios (though the differences were marginal in 2/8 scenarios), and only less stable in 1/8 scenarios (see Figures 5 and 6). Thus, even for the Shade and Stopnisek method, the branch-based approach appears to be advantageous.

The primary goal of this paper is to introduce a framework for phylogenetic analysis of core microbiomes. For this reason, we have focused on the concepts required to define a core community phylogeny and have outlined the application of a core community phylogeny to three types of microbiome analyses - Venn diagrams, ɑ-diversity (Faith’s PD), and β-diversity (UniFrac). Our broad approach, however, is not limited to these particular types of analyses. Indeed, additional analyses could be developed within the same framework. A straightforward example would be to use UniFrac distances between core microbiomes to project into ordination space (e.g., Principal Coordinate Analysis or Nonmetric Multidimensional Scaling). This would provide phylogenetically informed visualization of the relatedness of different core microbiomes.

While initially developed for microbiome analysis, beyond microbiomes, our community phylogenetics approach and associated R package may ultimately have wide application to macrobial systems as well. Currently, very few macrobial studies attempt to identify core communities (but see Murdock et al.^56^). This is likely because, at least historically, macrobial systems have not been studied with many replicates. Indeed, even when replicate macrobial communities are characterized, the effort required to identify all taxa in a single community means that replicate numbers are low. This makes it difficult to identify a core community. Further, in species poor macrobial systems where characterization of replicate communities is possible, the core community concept is not as necessary or useful because the systems are simple enough to analyze in their entirety. As sequencing techniques improve, however, and more and more macrobial systems are characterized in a manner similar to microbiomes (e.g., through the use of eDNA^57,58^), it seems likely that core communities will emerge as a common form of analysis for species rich macrobial systems (e.g., tropical rainforests or litter insects) as well. There is no reason why our proposed framework cannot be applied to macrobial communities. In fact, our framework can be applied to any system including prokaryotic and eukaryotic communities, host-associated and free-living communities, and microbial (virus, bacteria, fungi, etc.) and macrobial communities. It can also be applied to systems characterized through a wide variety of methods, even including non-sequencing techniques. The only requirements for our approach are tables of presences/absences of different taxa across different communities, along with a phylogenetic tree comprising all taxa present across all communities. Thus, occupancy data can be based on sequencing but can also be based on any other form of observation, including transect or plot surveys. Likewise, phylogenies can be constructed using sequence data, including sequencing of any kind (single marker genes, combinations of marker genes, whole genomes), but can also be generated based on morphology or other traits commonly used to build trees. While any phylogeny can be used for our phylogenetic analysis of core communities, the validity of the analysis will depend on the accuracy of the phylogeny; thus, care should be taken to select the most appropriate approach to tree building given the data available. For 16S rRNA gene sequencing, Qiime2 provides guidance and pipelines for both alignment-based and fragment-insertion methods of tree reconstruction.^44^ Likewise, for shotgun metagenomics sequencing, the Genome Taxonomy Database tookit (GTDB-Tk)^59,60^ provides different options for both *de novo* tree reconstruction and placement of genomes into a reference tree.

The identification and characterization of core microbiomes has become a hallmark of microbiome analysis. However, to date, most analyses of core microbiomes have focused on taxonomy at specified taxonomic ranks. By contrast, phylogeny-aware metrics for characterizing core microbiomes are largely lacking. Phylogeny-aware metrics have the benefit of integrating information across taxonomic ranks. This can be particularly helpful when analysis at different taxonomic ranks yields different outcomes - a common problem in ecology and evolution^61–65^ including the study of core microbiomes.^1^ Our phylogenetic framework overcomes this challenge. Indeed, even when our analysis is performed at different taxonomic ranks, it typically yields more consistent results than non-phylogenetic methods. With the development of our new framework and associated R package, we provide a mechanism to bring phylogeny-aware community analyses to the study of core microbiomes. Ultimately, this will allow for novel insight into the conserved components of microbial communities both within and between different types of habitats.

## Methods

### Datasets

#### Skin *Staphylococcus* microbiota

We downloaded the `ps_staph_ASV_filter_final.rds` file from https://github.com/skinmicrobiome/Joglekar_Staphylococcus_2023/tree/main/phyloseq_object. We then loaded this file into R and used it to write out a FASTA file of representative sequences. From the FASTA file, we used the `align-to-tree-mafft-fasttree` pipeline in Qiime2 to construct a phylogenetic tree for all *Staphylococcus* ASVs based on their 16S rRNA sequences. The FASTA file of representative sequences is available on our GitHub page. Next, we rarified samples to a read depth of 500 using the `rarefy_even_depth` function from the `phyloseq` package (version 1.50.0).^50^ For our analysis of individuals, we randomly selected 10 skin samples from all 9 individuals with 10+ skin samples available (recognizing that different body sites were used for different people). We then compared core microbiota across individuals based on ASVs and OTUs. For the latter, we agglomerated closely related taxa using the `tip_glom` function from the `phyloseq` package (h = 0.5). For our analysis of skin habitats, we randomly selected 20 skin samples from all 5 habitats with 20+ skin samples available (sebaceous, moist, dry, feet, nares). We performed all analyses of the *Staphylococcus* dataset using an unscaled phylogenetic tree. However, because many of the terminal branches were short, for the sake of visualization, we plotted the *Staphylococcus* phylogenies using a square-root transformation.

#### Lizard gut microbiota

We sampled the microbiota from several whiptail lizard species (genus *Aspidoscelis*, family *Teiidae*) at two locations (see Camper et al.^66^ for more site details) within Sevilleta National Wildlife Refuge. Each location represents a site of syntopy between hybrid *A. neomexicanus* and a progenitor species, *A. marmoratus* (♀; a heterogenous grassland-mesquite shrubland site: 34.397882°N, 106.867637°W, WGS84) and *A. arizonae* (♂, recently split from *A. inornatus*^67^; a grassland site with sparse cholla cacti: 34.331178°N, 106.636717°W, WGS84). Following taxonomic guidelines by Tucker et al.^68^ and Walker et al.,^69^ we refer to *Aspidoscelis* spp. using the masculine epithet. We captured lizards using a rapidly deployable drift fence trapping array lined with box funnel traps, further outlined in Camper et al.,^66^ and by hand or lizard noose when possible. Upon capture, we sampled the gut (=cloacal) and skin microbiota (for analysis of the skin dataset, see Additional file 1) of each species following a swabbing protocol in Camper et al.^22^ We sampled each location until we captured at least 15 female lizards of each whiptail species. All samples were sent to ZymoBIOMICS Targeted Sequencing Service for Microbiota Analysis (Zymo Research) using 16S rRNA gene sequencing. Additional details are available from https://www.zymoresearch.com/pages/16s-its-amplicon-sequencing and Camper et al.^22^ FASTQ files for lizard microbiomes, as well as environmental and sequencing controls, are available at (these will be uploaded to NCBI upon publication).

We used FASTA files of representative sequences, supplied to us by ZymioBIOMICS Targeted Sequencing Service (available on our GitHub page: https://github.com/bewicklab/CoreTrees/), to create phylogenetic trees, as described above. We then combined these trees with the .biom files, also supplied to us by the ZymoBIOMICS Targeted Sequencing Service (also available at: https://github.com/bewicklab/CoreTrees/) for all downstream analysis. Details on the Zymo pipeline can be found in our previous description of the dataset.^22^ Prior to analysis, we removed 16S rRNA gene sequences that could not be identified as Bacteria or Archaea based on the proprietary Zymo database. Additionally, we removed all samples with a read depth lower than 10,000 reads. Finally, we rarefied all samples to the read depth of the lowest sample. ^49^

#### Soil dataset

We downloaded dataset 1036 from redbiom (https://qiita.ucsd.edu/study/description/1036).^70^ We then used the `feature-table.qza` file and the `all.seqs.fa` files from the `130932` folder. From the FASTA file, we constructed a phylogeny, again using the same Qiime2 pipeline that was used for the *Staphylococcus* dataset. We then read the resulting feature table and phylogeny into R for analysis. For α-diversity, we rarefied samples to the size of the sample with the lowest read depth. For β-diversity, we rarefied samples to 5000 reads because of the computational time required by the `tip_glom` function for larger datasets (see below).

#### Lake dataset

We downloaded dataset 1041 from redbiom (https://qiita.ucsd.edu/study/description/1041). We then used the `feature-table.qza` file and the `reference-hit.seqs.fa` files from the `131527` folder. From the FASTA file, we constructed a phylogeny, again using the same Qiime2 pipeline that was used for the *Staphylococcus* dataset. We then read the resulting feature table and phylogeny into R for analysis. For both α-and β- diversity, we rarefied samples to the size of the sample with the lowest read depth. Analysis of the lake dataset can be found in Additional file 1.

### Dataset Analysis

#### Core Trees

We used the `coreEdges` function from our `holobiont` package to identify branches of the *Staphylococcus* phylogeny that were core to each individual. We then visualized the trees using a modification of the `plot_phylo` function from the `phyloseq` package (included in the code on our Github page). We used the `coreVennTree` function from our `holobiont` package to identify and visualize branches of the *Staphylococcus* phylogeny that were shared by the core microbiota of different skin habitats. For all figures in the main text, we used an occupancy threshold of 0.5 as our core criterion.

#### Venn Diagrams

We used the `coreVenn` and `corePhyloVenn` functions from our `holobiont` package to create non-phylogenetic and phylogenetic Venn Diagrams respectively. In all cases, we reported percentages of the total branch length and used rooted phylogenies. All Venn diagrams in the main text were calculated based on rooted trees for ASVs using an occupancy threshold of 0.5 as our core criterion. Additional core criteria are included in Additional file 1: Additional Venn Diagrams.

#### α-diversity

We used the `coreRichness` and `coreFaithsPD` functions from our `holobiont` package to calculate non-phylogenetic and phylogenetic α-diversity respectively. To compare across taxonomic ranks, we took the original ASV .biom file supplied by Zymo or .qza file downloaded from redbiom to generate an ASV phyloseq object.^50^ We then used the `tax_glom and tip_glom` functions from the `phyloseq` package to generate OTU tables and phylogenies for the lizard and soil/lake datasets respectively. For `tax_glom,` we considered taxonomic ranks from species (‘Rank7’) through to order (‘Rank4’). For `tip_glom` we considered cophenetic distances from h = 0.05 to h = 0.2. This allowed us to calculate Faith’s PD at different taxonomic ranks/levels of tip agglomeration, as well as related non-phylogenetic richness. All Faith’s PD and richness calculations were performed on rooted trees using the following core criteria: (1) an occupancy threshold of 0.5, (2) an occupancy threshold of 0.5 and a mean relative abundance threshold of 0.1% and (3) a modified Shade and Stopnisek method (see Additional File 1: Shade and Stopnisek Algorithm).

#### β-diversity

We used the `coreJaccard` and `coreUniFrac` functions from our `holobiont` package to calculate non-phylogenetic and phylogenetic β-diversity respectively. Comparisons across taxonomic ranks/tip agglomeration were performed as described for Faith’s PD. Again, all UniFrac and Jaccard calculations were performed on rooted trees using the following core criteria: (1) an occupancy threshold of 0.5, (2) an occupancy threshold of 0.5 and a mean relative abundance threshold of 0.1% and (3) a modified Shade and Stopnisek method.

All analyses were performed using the R programming language^71^ (version 4.2.1) available on our Github page at: https://github.com/bewicklab/CoreTrees

## Supporting information

BIOM file

## Declarations

### Ethics approval and consent to participate

All research was approved by Clemson University under IACUC protocol numbers #2020-015 and #2021-047. We completed this work under the Sevilleta National Wildlife Refuge Special Use Permit #SEV_Bewick_Camper_2022_59 and the New Mexico Department of Game and Fish permit authorization #3772.

### Consent for publication

not applicable

### Availability of data and materials

All data and code (BIOM files, metadata files, and all R code) necessary for the analyses presented in this manuscript are available at https://github.com/bewicklab/CoreTrees unless otherwise noted in the main text. All data (BIOM and metadata files) are further included as a zipped file as Additional file 2. The `holobiont` R package is publicly available through CRAN, available at https://cran.r-project.org/web/packages/holobiont/index.html. We have uploaded all raw sequence data (lizard microbiota and controls) to the National Center for Biotechnology Information’s SRA database (https://dataview.ncbi.nlm.nih.gov/object/PRJNA1188019?reviewer=9hrl0g7nl4fmdqf8icgjglbes <u>2</u>), and this information will be made publicly available upon acceptance.

### Authors’ contributions

BTC and SAB conceived the idea and wrote the first draft of the manuscript. BTC performed all related fieldwork. SAB developed the code and data analysis. All authors contributed substantially to the content of the manuscript.

## Acknowledgements

We thank Thomas Dempster, August Spencer, Eva Purcell, Riley Manuel, Drew Kanes and Zachary Laughlin for their field assistance in New Mexico as well as Lily Margeson, Simon Dunn, Georgianna Bellinger, Henry Egloff, Kaila Hodges, Camryn Lachica, and Savannah Utz for their assistance assembling drift fence trapping arrays.

## Electronic Supplementary Material

core_community_phylogeny_example.docx

Additional file 1: A simplified system demonstrating the construction of core community phylogenies and calculations of associated metrics.

biom_files.zip

Additional file 2: BIOM files, read counts, taxonomy assignments, microbial phylogenies, and lizard metadata for gut and skin samples used in this study.

