## Additional file 1 for "Phylogenetic Measures of the Core Microbiome"

### 1. Simple Example System:

Consider three different habitats (red, blue, and green) and 10 microbial taxa (A-J). Tables S1.1-S1.3 show presence/absence (P/A) data for microbiome samples from each habitat.

**Table S1.1.** P/A data for 16 samples from the red habitat; the final column shows the total number of samples from the red habitat that each taxon was found in.

| Taxon | R1 | R2 | R3 | R4 | R5 | R6 | R7 | R8 | R9 | R10 | R11 | R12 | R13 | R14 | R15 | R16 | Rtotal |
|---|---|---|---|---|---|---|---|---|---|---|---|---|---|---|---|---|---|
| A | 0 | 1 | 0 | 1 | 0 | 1 | 0 | 0 | 0 | 0 | 0 | 0 | 0 | 0 | 0 | 0 | 3 |
| B | 0 | 0 | 0 | 1 | 0 | 0 | 0 | 1 | 0 | 0 | 0 | 0 | 0 | 0 | 0 | 1 | 3 |
| C | 0 | 1 | 0 | 0 | 0 | 1 | 0 | 1 | 1 | 0 | 0 | 0 | 1 | 0 | 0 | 1 | 6 |
| D | 0 | 1 | 0 | 1 | 1 | 0 | 0 | 0 | 1 | 1 | 0 | 1 | 1 | 0 | 0 | 0 | 7 |
| E | 1 | 1 | 1 | 1 | 0 | 0 | 0 | 1 | 1 | 1 | 0 | 1 | 1 | 0 | 1 | 1 | 11 |
| F | 0 | 0 | 0 | 1 | 0 | 0 | 0 | 1 | 0 | 0 | 0 | 0 | 0 | 0 | 0 | 0 | 2 |
| G | 1 | 1 | 0 | 1 | 0 | 0 | 0 | 0 | 1 | 0 | 1 | 1 | 1 | 0 | 1 | 0 | 8 |
| H | 0 | 0 | 0 | 0 | 0 | 0 | 0 | 0 | 0 | 0 | 0 | 1 | 0 | 0 | 0 | 0 | 1 |
| I | 1 | 1 | 1 | 1 | 1 | 1 | 1 | 1 | 1 | 0 | 1 | 1 | 1 | 1 | 0 | 1 | 14 |
| J | 1 | 1 | 0 | 0 | 0 | 0 | 0 | 0 | 0 | 0 | 1 | 1 | 1 | 0 | 1 | 1 | 7 |

**Table S1.2.** P/A data for 15 samples from the blue habitat; the final column shows the total number of samples from the blue habitat that each taxon was found in.

| Taxon | B1 | B2 | B3 | B4 | B5 | B6 | B7 | B8 | B9 | B10 | B11 | B12 | B13 | B14 | B15 | Btotal |
|---|---|---|---|---|---|---|---|---|---|---|---|---|---|---|---|---|
| A | 0 | 0 | 0 | 1 | 0 | 0 | 0 | 0 | 0 | 0 | 0 | 1 | 1 | 0 | 1 | 4 |
| B | 0 | 0 | 0 | 1 | 0 | 0 | 0 | 0 | 0 | 0 | 0 | 1 | 1 | 0 | 0 | 3 |
| C | 0 | 0 | 0 | 0 | 0 | 0 | 0 | 0 | 0 | 0 | 0 | 0 | 1 | 0 | 0 | 1 |
| D | 0 | 0 | 0 | 0 | 1 | 0 | 1 | 0 | 0 | 0 | 0 | 1 | 1 | 1 | 1 | 6 |
| E | 0 | 1 | 1 | 0 | 1 | 1 | 1 | 0 | 0 | 1 | 0 | 1 | 1 | 1 | 1 | 10 |
| F | 0 | 0 | 0 | 0 | 1 | 0 | 0 | 0 | 0 | 0 | 0 | 1 | 1 | 1 | 1 | 5 |
| G | 0 | 0 | 0 | 0 | 0 | 0 | 0 | 0 | 0 | 0 | 0 | 1 | 0 | 1 | 0 | 2 |
| H | 0 | 0 | 1 | 1 | 0 | 0 | 0 | 1 | 0 | 0 | 0 | 0 | 1 | 0 | 0 | 4 |
| I | 0 | 1 | 1 | 1 | 1 | 1 | 1 | 1 | 1 | 1 | 1 | 1 | 1 | 1 | 1 | 14 |
| J | 1 | 0 | 0 | 1 | 0 | 0 | 0 | 0 | 0 | 1 | 0 | 1 | 1 | 1 | 0 | 6 |

**Table S1.3.** P/A data for 34 samples from the green habitat; the final column shows the total number of samples from the green habitat that each taxon was found in.

| Taxon | G1 | G2 | G3 | G4 | G5 | G6 | G7 | G8 | G9 | G10 | G11 | G12 | G13 | G14 | G15 | G16 | G17 | G18 | G19 | G20 | G21 | G22 | G23 | G24 | G25 | G26 | G27 | G28 | G29 | G30 | G31 | G32 | G33 | G34 | Gtotal |
|---|---|---|---|---|---|---|---|---|---|---|---|---|---|---|---|---|---|---|---|---|---|---|---|---|---|---|---|---|---|---|---|---|---|---|---|
| A | 0 | 0 | 0 | 0 | 0 | 0 | 0 | 1 | 0 | 1 | 0 | 0 | 0 | 1 | 0 | 0 | 0 | 0 | 0 | 0 | 0 | 0 | 0 | 0 | 1 | 0 | 0 | 0 | 0 | 0 | 1 | 0 | 0 | 1 | 6 |
| B | 0 | 0 | 0 | 1 | 0 | 0 | 1 | 1 | 0 | 1 | 0 | 1 | 0 | 1 | 1 | 1 | 0 | 1 | 1 | 0 | 0 | 0 | 0 | 1 | 1 | 1 | 1 | 1 | 0 | 1 | 0 | 1 | 1 | 0 | 18 |
| C | 1 | 0 | 0 | 0 | 0 | 0 | 0 | 0 | 0 | 1 | 0 | 0 | 0 | 0 | 1 | 1 | 0 | 1 | 0 | 1 | 1 | 1 | 0 | 0 | 1 | 1 | 0 | 1 | 1 | 0 | 1 | 0 | 0 | 0 | 13 |
| D | 0 | 1 | 1 | 1 | 0 | 0 | 1 | 0 | 0 | 1 | 0 | 1 | 0 | 0 | 0 | 1 | 0 | 0 | 1 | 0 | 0 | 0 | 0 | 0 | 0 | 0 | 1 | 0 | 1 | 1 | 0 | 1 | 0 | 0 | 12 |
| E | 1 | 1 | 1 | 1 | 1 | 1 | 1 | 1 | 1 | 0 | 1 | 1 | 1 | 1 | 1 | 1 | 0 | 1 | 1 | 1 | 1 | 1 | 1 | 1 | 1 | 1 | 1 | 1 | 1 | 1 | 1 | 1 | 1 | 1 | 32 |
| F | 0 | 1 | 1 | 0 | 1 | 0 | 0 | 1 | 0 | 0 | 1 | 0 | 0 | 1 | 1 | 1 | 0 | 1 | 1 | 0 | 0 | 1 | 0 | 0 | 0 | 0 | 1 | 1 | 0 | 0 | 0 | 1 | 0 | 1 | 15 |
| G | 0 | 0 | 0 | 0 | 0 | 0 | 0 | 0 | 0 | 1 | 0 | 0 | 0 | 0 | 0 | 0 | 1 | 1 | 0 | 0 | 0 | 0 | 0 | 0 | 0 | 0 | 0 | 0 | 0 | 1 | 0 | 0 | 1 | 0 | 5 |
| H | 0 | 0 | 0 | 0 | 0 | 0 | 1 | 1 | 0 | 1 | 1 | 0 | 0 | 0 | 0 | 0 | 0 | 0 | 0 | 0 | 0 | 0 | 0 | 0 | 0 | 0 | 0 | 0 | 1 | 0 | 0 | 0 | 0 | 1 | 6 |
| I | 1 | 1 | 1 | 1 | 1 | 0 | 1 | 1 | 1 | 1 | 1 | 1 | 1 | 1 | 1 | 1 | 1 | 1 | 1 | 1 | 1 | 1 | 1 | 1 | 1 | 1 | 1 | 1 | 1 | 1 | 1 | 1 | 1 | 1 | 33 |
| J | 0 | 1 | 0 | 0 | 0 | 1 | 1 | 1 | 1 | 1 | 1 | 0 | 1 | 1 | 0 | 0 | 0 | 0 | 1 | 1 | 0 | 1 | 1 | 1 | 0 | 1 | 1 | 0 | 0 | 0 | 0 | 0 | 0 | 1 | 17 |

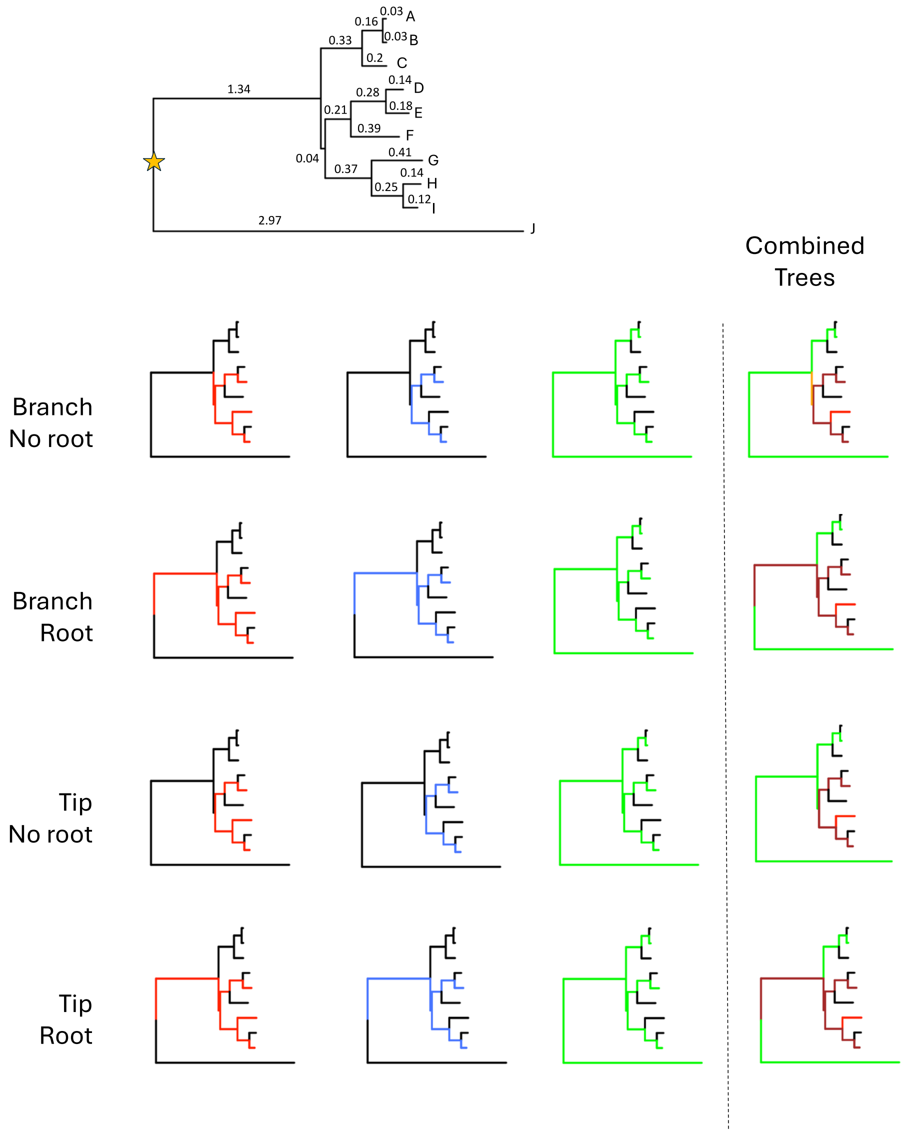


**Figure S1.1.** Phylogenies for our simplified example (Tables S1.1-S1.3). The top panel shows the overall phylogenetic tree for microbial taxa A-J. Branch lengths are labelled on individual branches and a star is used to identify the root node/MRCA. The first, second, and third columns in the lower panels show the core community phylogenies for the three different types of habitats (red, blue, green). The fourth column shows which branches are shared by core communities from which combinations of habitats as follows: only core in the first habitat (red), only core in the second habitat (blue), only core in the third habitat (green), only core in the first and second habitats (purple), only core in the first and third habitats (orange), only core in the second and third habitats (turquoise), core in all three habitats (brown), and core in no habitats (black). The first row shows branch-based phylogenies assuming an unrooted tree, the second row shows branch-based phylogenies assuming a rooted tree, the third row shows tip-based phylogenies assuming an unrooted tree, and the fourth row shows tip-based phylogenies assuming an unrooted tree.

***Example of Core Phylogenies and Venn Diagrams:***

Figure S1.1 shows the phylogeny for our simplified system, as well as the core community phylogenies for the red, blue, and green habitats assuming a core threshold of 50% (i.e., the taxon or branch must occur in at least 8 samples from the red and blue habitats and in at least 17 samples from the green habitat). Figure S1.1 also shows the combined core community phylogenies colored based on the branches that are shared by different combinations of the three different habitats. Notice that, when using an unrooted tree, the branch basal to the D-J clade is included in the branch-based phylogeny for the red community but not for the blue community. This highlights the challenge of constructing branch-based phylogenies on unrooted trees. In the red community, at least one member of the D-J clade appears in the same sample/community as at least one member of the A-C clade or species J in 11 of the 16 samples (R1, R2, R4, R6, R8, R9, R11, R12, R13, R15, and R16). Thus, the branch basal to the D-J clade surpasses the 50% threshold. However, in the blue community, at least one member of the D-J clade appears in the same sample as at least one member of the A-C clade or species J in only 6 of the 15 samples (B4, B10, B12, B13, B14, B15). Thus, the branch basal to the D-J clade does not surpass the 50% threshold. This dependence of the unrooted, branch-based core community phylogeny, not only on overall taxon occurrence rates, but also on taxon co-occurrence patterns across samples is why we suggest always using a rooted tree. When using a rooted tree, all samples necessarily include the root node, removing dependence on co-occurrence of taxa across samples. That said, because most individual microbiome samples contain multiple microbial phyla, and thus span deeply divergent clades, rooted versus unrooted trees will likely yield nearly identical results. Indeed, the type of anomalous behavior that might arise from using unrooted trees is more likely to be problematic in macrobial systems, where both lower diversity and narrower taxonomic focus may result in individual communities that do not span most of the phylogeny of the entire habitat or set of habitats.

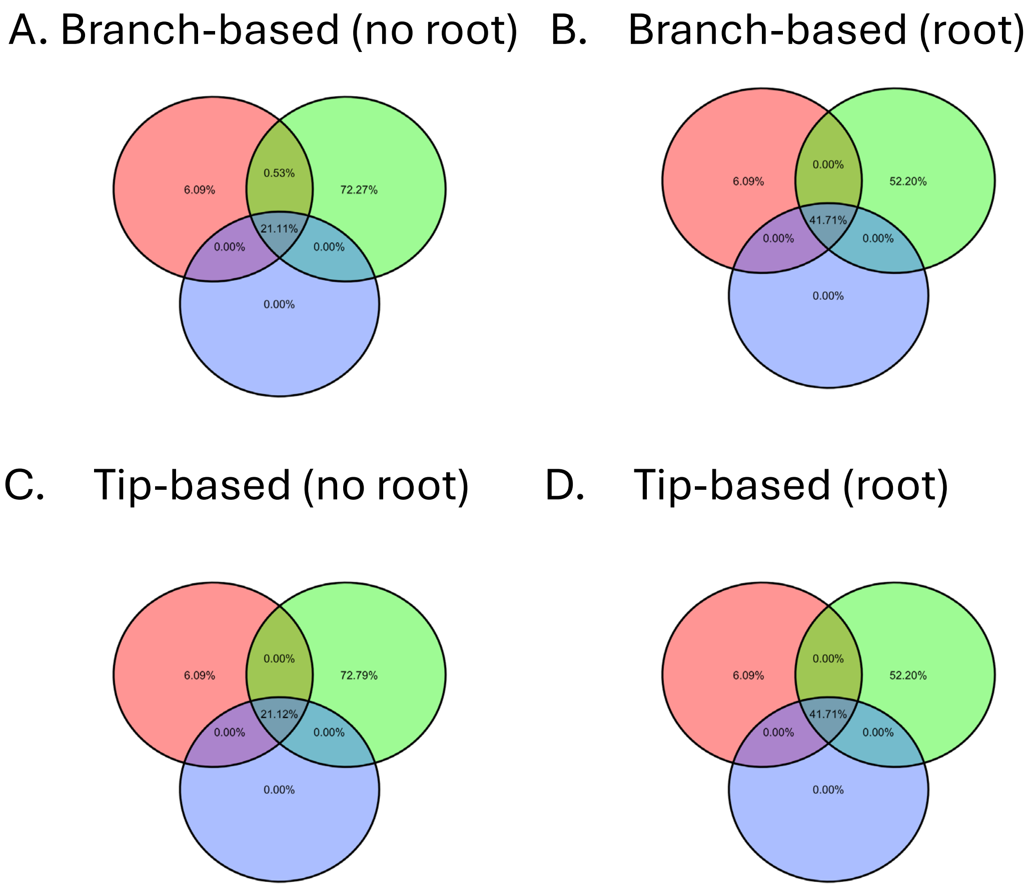


**Figure S1.2.** Venn diagrams for each of the possible core community phylogenies using the same communities as in Figure S1.1.

Figure S1.2 shows phylogenetic Venn diagrams for the red, blue and green habitats using each of our four methods (rooted and unrooted tip and branch-based approaches). While results are largely consistent, there are slight differences based on which internal branches are included. Thus, both non-rooted methods deviate from their corresponding rooted method, while the non-rooted tip-based and branch-based methods differ slightly from one another. Again, unless there is strong reason to do so, we suggest using the rooted branch-based method. Nevertheless, our R package provides functionality for all three options.

#### *Example Calculations of Faith’s PD:*

Core Faith’s PD can be calculated by summing the total branch length in a particular core community phylogeny. Below we explicitly show the branch length summations for two core community phylogenies – one using the unrooted, branch-based approach, and the other using the rooted, tip-based approach. Table S1.4 provides the values of core Faith’s PD for all possible methods for each of our simple example systems.

$$\mathrm{PD}_{\mathrm{Red,branch,no\ root}} = 0.12 + 0.25 + 0.41 + 0.37 + 0.04 + 0.21 + 0.28 + 0.18 = 1.86$$

$$\mathrm{PD}_{\mathrm{Blue,tip,root}} = 1.34 + 0.04 + 0.21 + 0.37 + 0.28 + 0.18 + 0.25 + 0.12 = 2.79$$

| | Red | Green | Blue |
|---|---|---|---|
| [illegible] | 1.854828 | 6.283959 | 1.412300 |
| [illegible] | 3.197638 | 6.283959 | 2.790331 |
| [illegible] | 1.819607 | 6.283959 | 1.412300 |
| Tip-based, root | 3.197638 | 6.283959 | 2.790331 |

***Example Calculations of UniFrac Distance:***

The core UniFrac distance can be calculated by dividing the length of the unique branches of each core community phylogeny by the total length of all the branches in all core community phylogenies. Below we explicitly show this quotient for two core community phylogenies – one using the unrooted, branch-based approach, and the other using the rooted, tip-based approach. Table S1.5 provides the values of core UniFrac distances for all possible methods for each of our simple example systems.

$$\mathrm{UniFrac}_{\mathrm{Red|Blue,branch,no\ root}} = \frac{0.41 + 0.04}{0.12 + 0.25 + 0.41 + 0.37 + 0.04 + 0.21 + 0.28 + 0.18} = 0.24$$

$$\mathrm{UniFrac}_{\mathrm{Blue|Green,tip,root}}$$

$$= \frac{0.03 + 0.16 + 0.33 + 2.97}{0.03 + 0.16 + 0.33 + 1.34 + 0.04 + 2.97 + 0.12 + 0.25 + 0.37 + 0.21 + 0.28 + 0.18} = 0.56$$

**Table S1.5.** UniFrac distance between each pair of habitats from Figure S1.1.

| | Community | | |
|---|---|---|---|
| | Red/Blue | Red/Green | Green/Blue |
| Branch-based, no root | 0.238582 | 0.78367 | 0.775253 |
| Branch-based, root | 0.127378 | 0.582989 | 0.55596 |
| Tip-based, no root | 0.223844 | 0.788934 | 0.775253 |
| Tip-based, root | 0.127378 | 0.582989 | 0.55596 |

### 2. Additional *Staphylococcus* Analyses

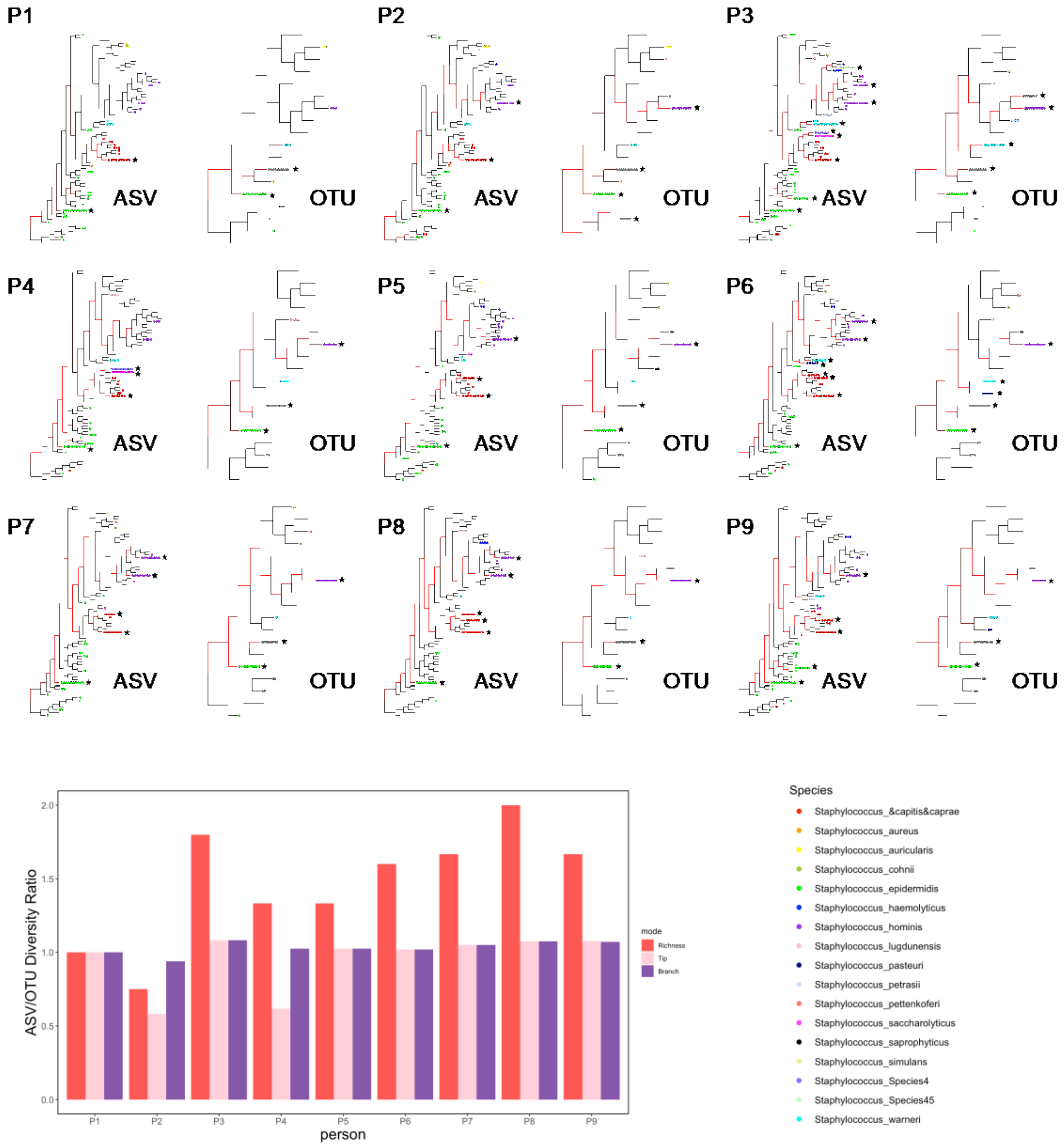


**Figure S2.1** (top) *Staphylococcus* phylogenies for 10 skin microbiome samples from 9 individuals using ASVs (left) and OTUs (right). Circles at the end of each terminal branch indicate the number of samples that contained the corresponding ASV or OTU. These circles are colored based on assigned species, with grey being used when multiple species are contained within the same OTU. Black stars indicate taxa that would be part of the core microbiome based on a non-phylogenetic approach using an occupancy threshold of 50%. Red lines indicate branches that are part of the core microbiome based on a branch-based phylogenetic approach using the same occupancy threshold of 50%. (bottom) ASV:OTU core *Staphylococcus* diversity for each of the 9 individuals using richness (red), tip-based (pink) and branch-based (purple) Faith's PD.

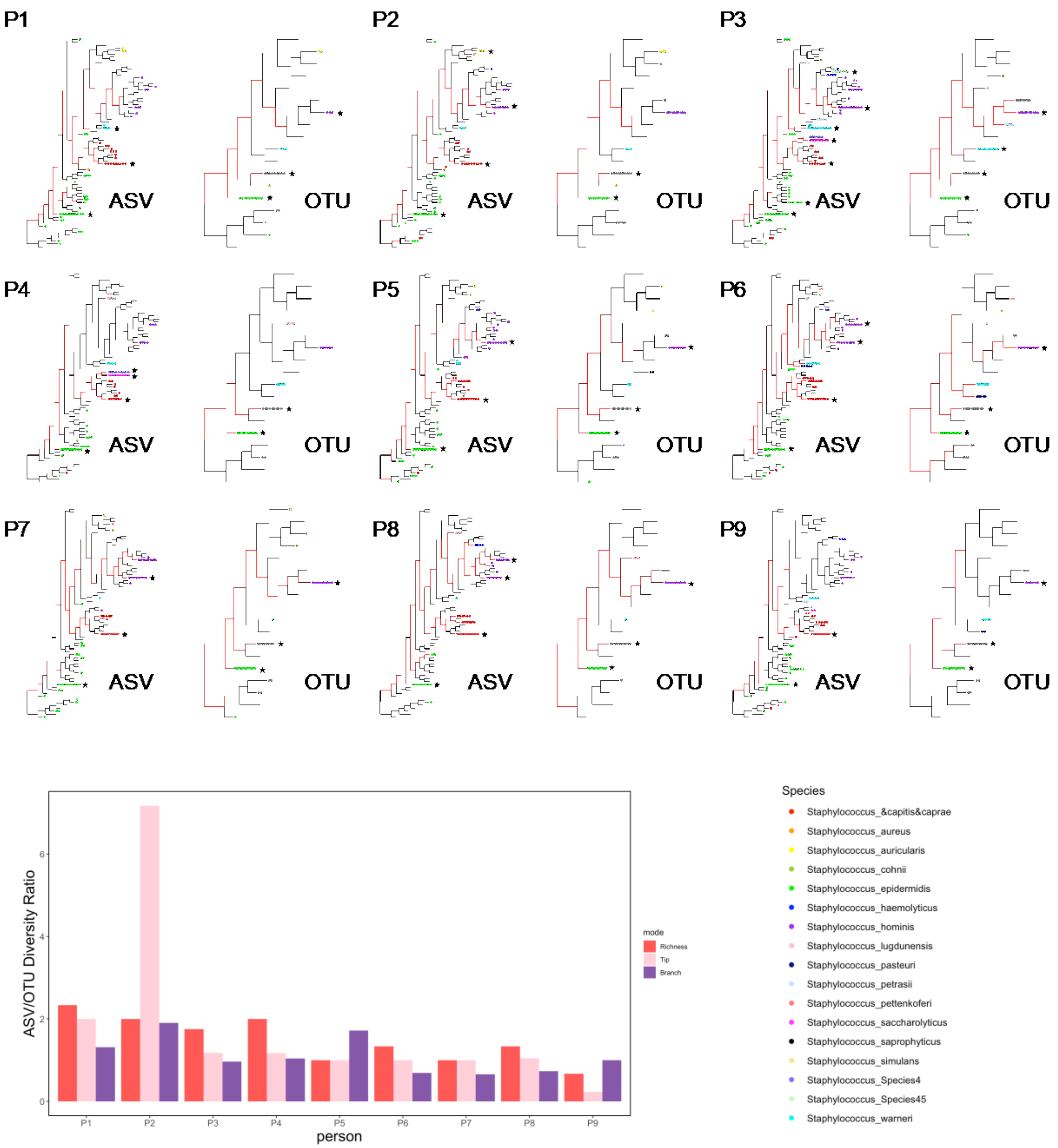


**Figure S2.2** (top) *Staphylococcus* phylogenies for 10 skin microbiome samples from 9 individuals using ASVs (left) and OTUs (right). Circles at the end of each terminal branch indicate the number of samples that contained the corresponding ASV or OTU. These circles are colored based on assigned species, with grey being used when multiple species are contained within the same OTU. Black stars indicate taxa that would be part of the core microbiome based on a non-phylogenetic approach using the Shade and Stopnisek method. Red lines indicate branches that are part of the core microbiome based on a branch-based phylogenetic approach using the same Shade and Stopnisek method. (bottom) ASV:OTU core *Staphylococcus* diversity for each of the 9 individuals using richness (red), tip-based (pink) and branch-based (purple) Faith's PD.

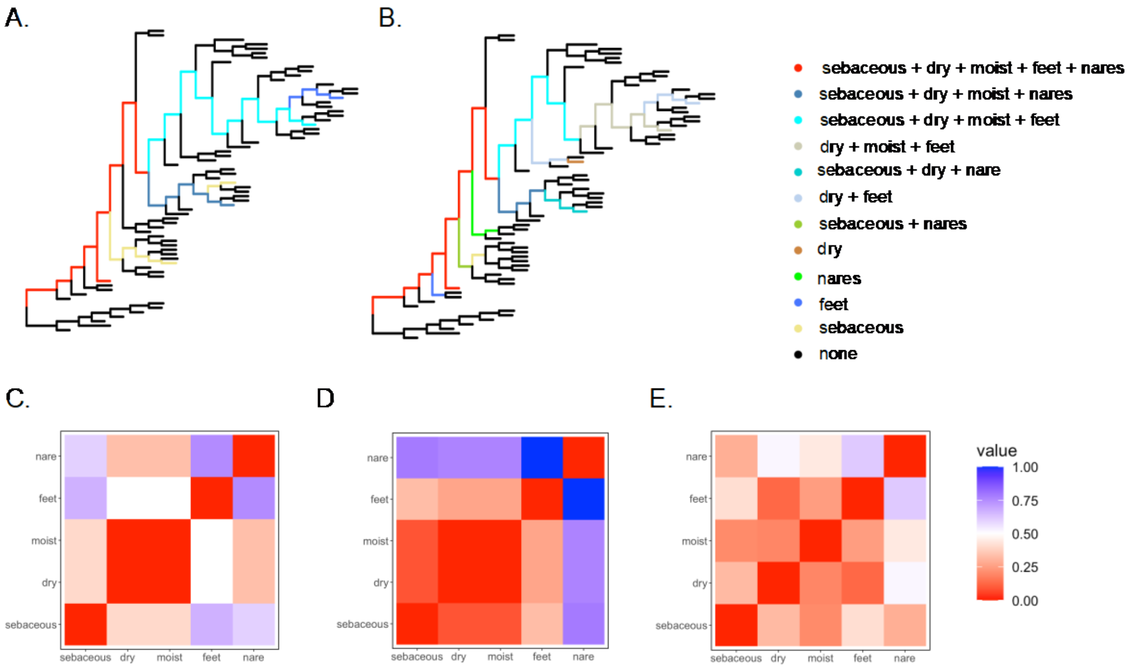


**Figure S2.3** (A) Tip-based and (B) branch-based *Staphylococcus* phylogenies showing the shared and unique core community phylogenies, along with Jaccard distances (C), tip-based UniFrac distances (D) and branch-based UniFrac distances (E) among skin habitats. All panels in this figure used the Shade and Stopnisek core criterion.

### 3. Additional Venn Diagrams

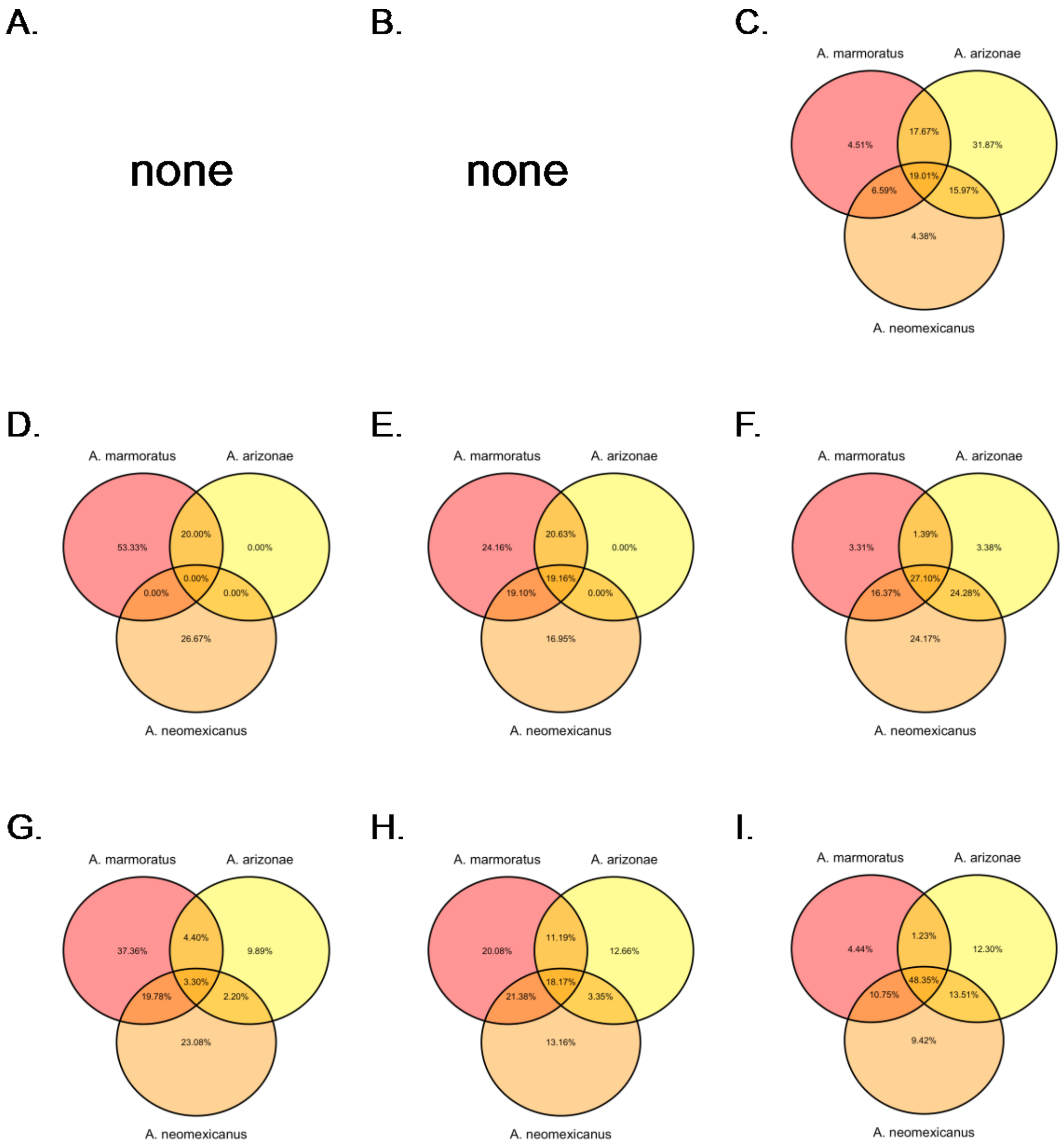


**Figure S3.1** Venn diagrams for lizard gut microbiota showing the percentage of the total ASV counts (A,D,G) and the phylogenetic branch length shared by the core microbiota from each lizard species using either tip-based (B,E,H) or branch-based (C,F,I) phylogenies. In (A-C) we use a 99% occupancy threshold as the core inclusion criterion. In (D-F) we use a 50% occupancy threshold and a 0.1% mean abundance threshold as the core inclusion criterion. In (G-I) we use the Shade and Stopnisek method as the core inclusion criterion. In all Venn diagrams, lizard populations are as follows: *Aspidoscelis arizonae* (yellow), *A. neomexicanus* (orange), *A. marmoratus* (red).

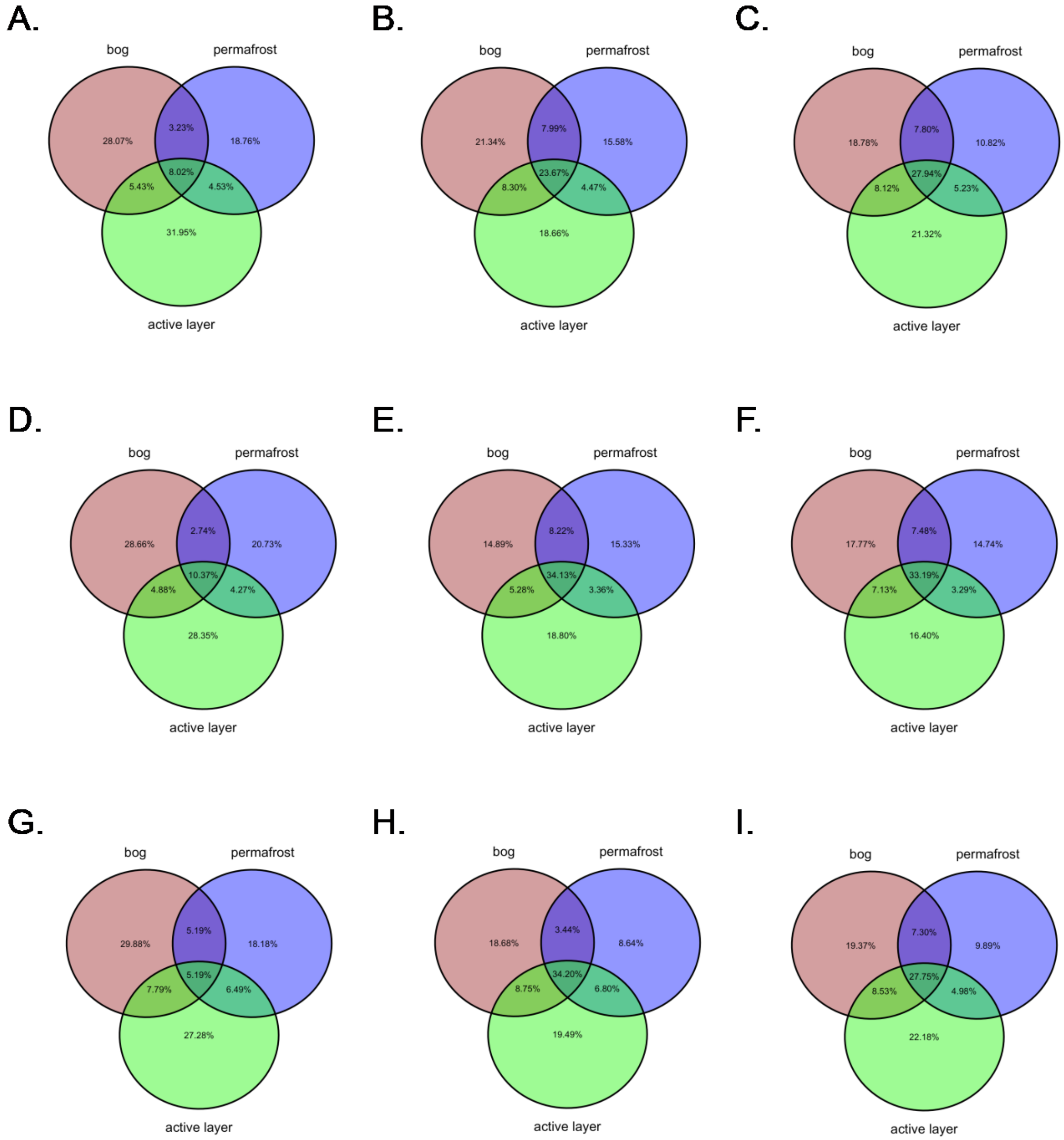


**Figure S3.2** Venn diagrams for soil microbiota showing the percentage of the total ASV counts (A,D,G) and the phylogenetic branch length shared by the core microbiota from each soil type using either tip-based (B,E,H) or branch-based (C,F,I) phylogenies. In (A-C) we use a 99% occupancy threshold as the core inclusion criterion. In (D-F) we use a 50% occupancy threshold and a 0.1% mean abundance threshold as the core inclusion criterion. In (G-I) we use the Shade and Stopnisek method as the core inclusion criterion. In all Venn diagrams, soil types are as follows: bog (brown), active layer (green), permafrost (blue).

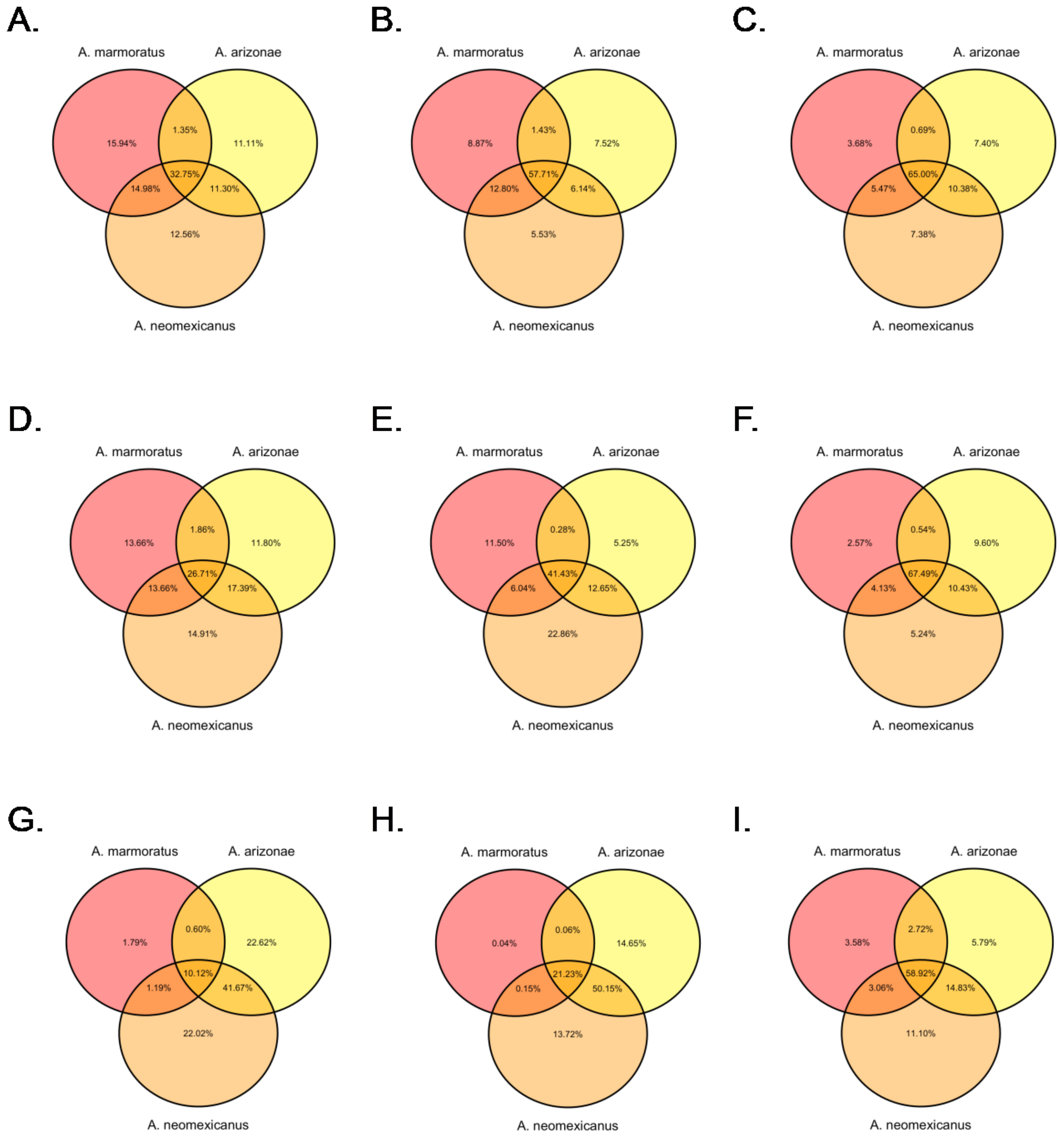


**Figure S3.3** Venn diagrams for lizard skin microbiota showing the percentage of the total ASV counts (A,D,G) and the phylogenetic branch length shared by the core microbiota from each lizard species using either tip-based (B,E,H) or branch-based (C,F,I) phylogenies. In (A-C) we use a 50% occupancy threshold as the core inclusion criterion. In (D-F) we use a 50% occupancy threshold and a 0.1% mean abundance threshold as the core inclusion criterion. In (G-I) we use the Shade and Stopnisek method as the core inclusion criterion. In all Venn diagrams, lizard populations are as follows: *Aspidoscelis arizonae* (yellow), *A. neomexicanus* (orange), *A. marmoratus* (red).

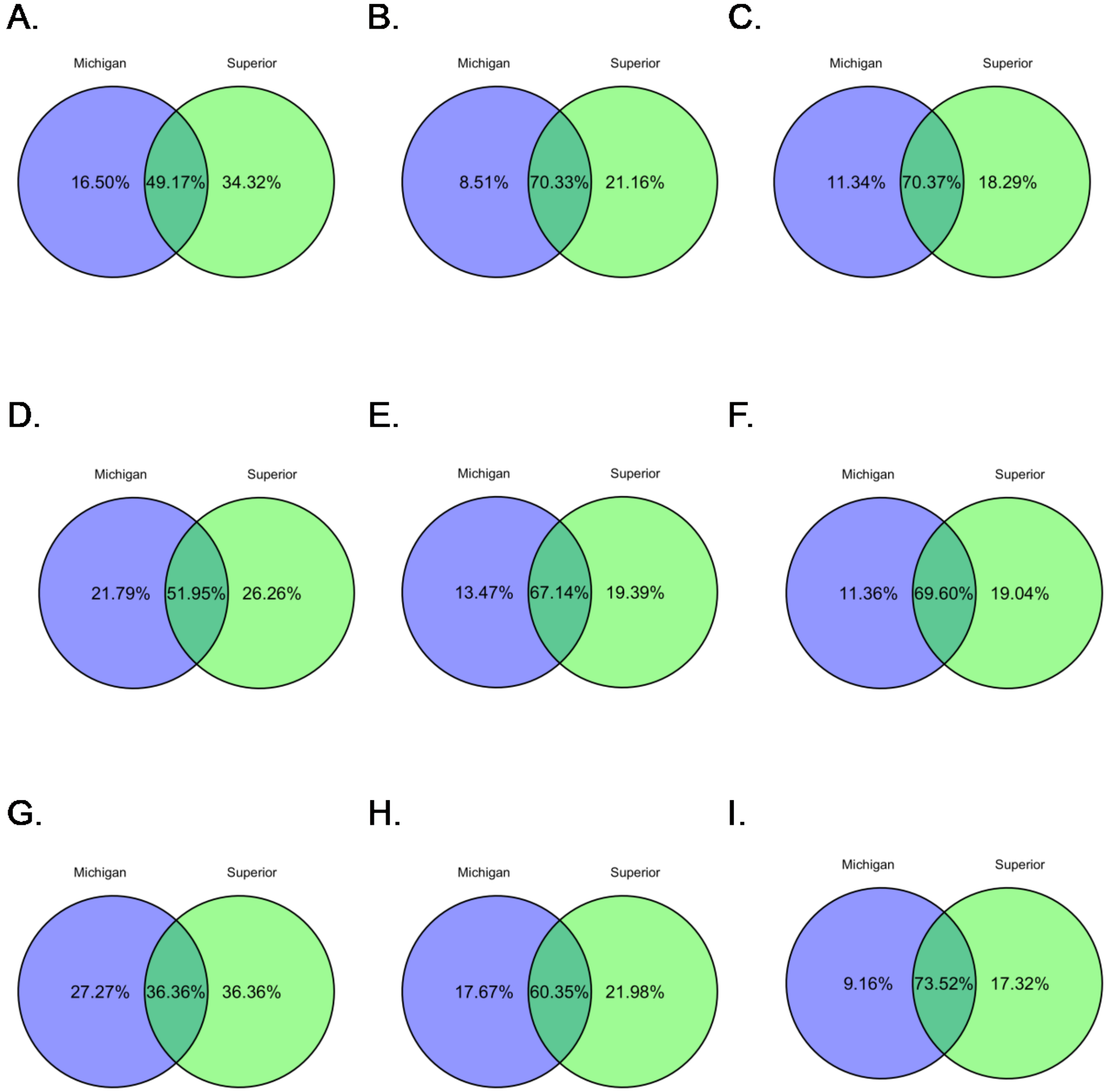


**Figure S3.4** Venn diagrams for lake water microbiota showing the percentage of the total ASV counts (A,D,G) and the phylogenetic branch length shared by the core microbiota from Lakes Michigan and Superior using either tip-based (B,E,H) or branch-based (C,F,I) phylogenies. In (A-C) we use a 50% occupancy threshold as the core inclusion criterion. In (D-F) we use a 50% occupancy threshold and a 0.1% mean abundance threshold as the core inclusion criterion. In (G-I) we use the Shade and Stopnisek method as the core inclusion criterion. In all Venn diagrams, lake microbiota are as follows: Lake Michigan (blue), Lake Superior (green).

### 4. Additional α- and β-Diversity Analyses

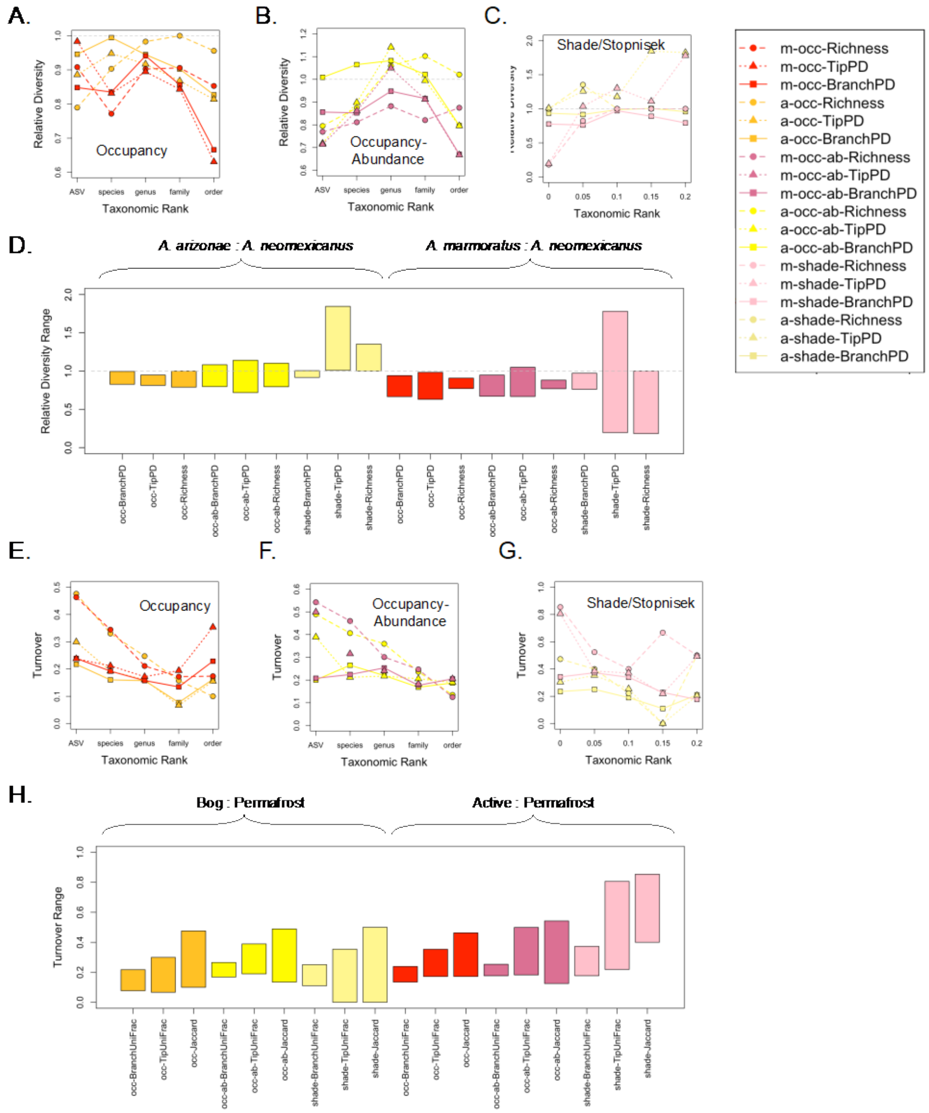


**Figure S4.1.** Comparison of α- (A-D) and β- (E-H) diversity of *Aspidoscelis* skin microbiota across taxonomic ranks for the branch-based Core Faith's PD (solid, squares), tip-based Core Faith's PD (dotted, triangles), and richness (dashed, circles) assuming the following core criteria: (A,E) an occupancy threshold of 50% (gold, red), (B,F) an occupancy threshold of 50% and a mean abundance threshold of 0.1% (yellow, dark pink), (C,G) the Shade and Stopnisek method (light yellow, light pink). In all panels we show *Aspidoscelis arizonae* (gold, yellow, pale yellow) and *A. marmoratus* (red, dark pink, light pink) relative/compared to *A. neomexicanus*. Diversity metrics are indicated as follows: 'BranchPD' is the branch-based phylogeny approach,

'TipPD' is the tip-based phylogeny approach, and 'Richness' is a count of taxa for each taxonomic rank. Core criteria are indicated as follows: 'occ' is the 50% occupancy threshold, 'occ-ab' is the 50% occupancy and 0.1% abundance threshold, and 'shade' is the Shade and Stopnisek method.

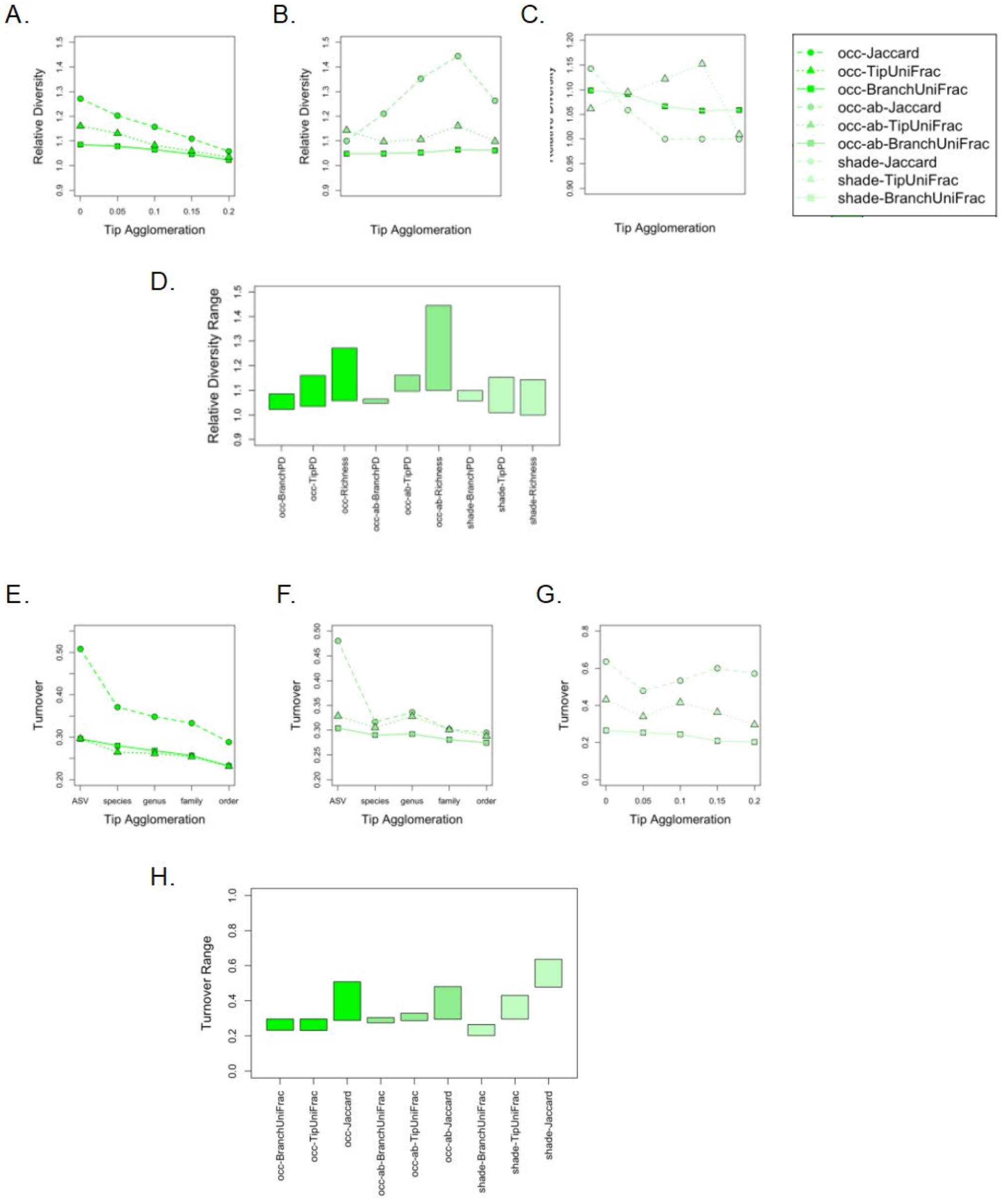


**Figure S4.2.** Comparison of α- (A-D) and β- (E-H) diversity of lake microbiota as a function of tip agglomeration for the branch-based Core Faith's PD (solid, squares), tip-based Core Faith's PD (dotted, triangles), and richness (dashed, circles) assuming the following core criteria: (A,E) an occupancy threshold of 50% (bright green), (B,F) an occupancy threshold of 50% and a mean abundance threshold of 0.1% (medium green), (C,G) the Shade and Stopnisek method (light green). In all panels we show Lake Superior relative/compared to Lake Michigan. Diversity metrics are indicated as follows: ‘BranchPD’ is the branch-based phylogeny approach, ‘TipPD’ is the tip-based phylogeny approach, and ‘Richness’ is a count of taxa for each taxonomic rank. Core criteria are indicated as follows: ‘occ’ is the 50% occupancy threshold, ‘occ-ab’ is the 50% occupancy and 0.1% abundance threshold, and ‘shade’ is the Shade and Stopnisek method.

**5. Shade and Stopnisek Algorithm**

The holobiont package includes a modified version of the Shade and Stopnisek method for identifying core microbiota. Briefly, we begin by ranking taxa or branches based on occupancy. Within each occupancy class, we then rank taxa or branches based on total abundance. Using the ranked occupancy-abundance list, we begin with the first taxon (tip-based) or the first set of branches (branch-based, the default is to begin with all branches present in every sample), and consecutively add taxa/branches. Each time we add a taxon/branch, we calculate the contributions of that taxon/branch to Bray-Curtis (tip-based) or weighted Unifrac (branch-based) distances between samples. We do this up to a maximum number of taxa/branches (the default is 1% of taxa/branches). We then divide the Bray-Curtis/weighted Unifrac contributions from any given taxon/branch by the total Bray-Curtis/weighted Unifrac distances for all taxa/branches. This gives percent contributions of each taxon/branch. Next, we calculate the mean percent contribution of each taxon/branch across all pairs of samples. Finally, we find the lowest occupancy-abundance ranked taxon/branch (i.e., lowest occupancy/abundance) that yields a threshold percent increase (the default is 2%) in Bray-Curtis/weighted Unifrac distance. The core microbiota is then all taxa/branches with higher occupancy/abundance than this final taxon. This algorithm is implemented in the `shade_np`, `shade_tip` and `shade_branch` functions of the holobiont package. R code is available from our GitHub page.

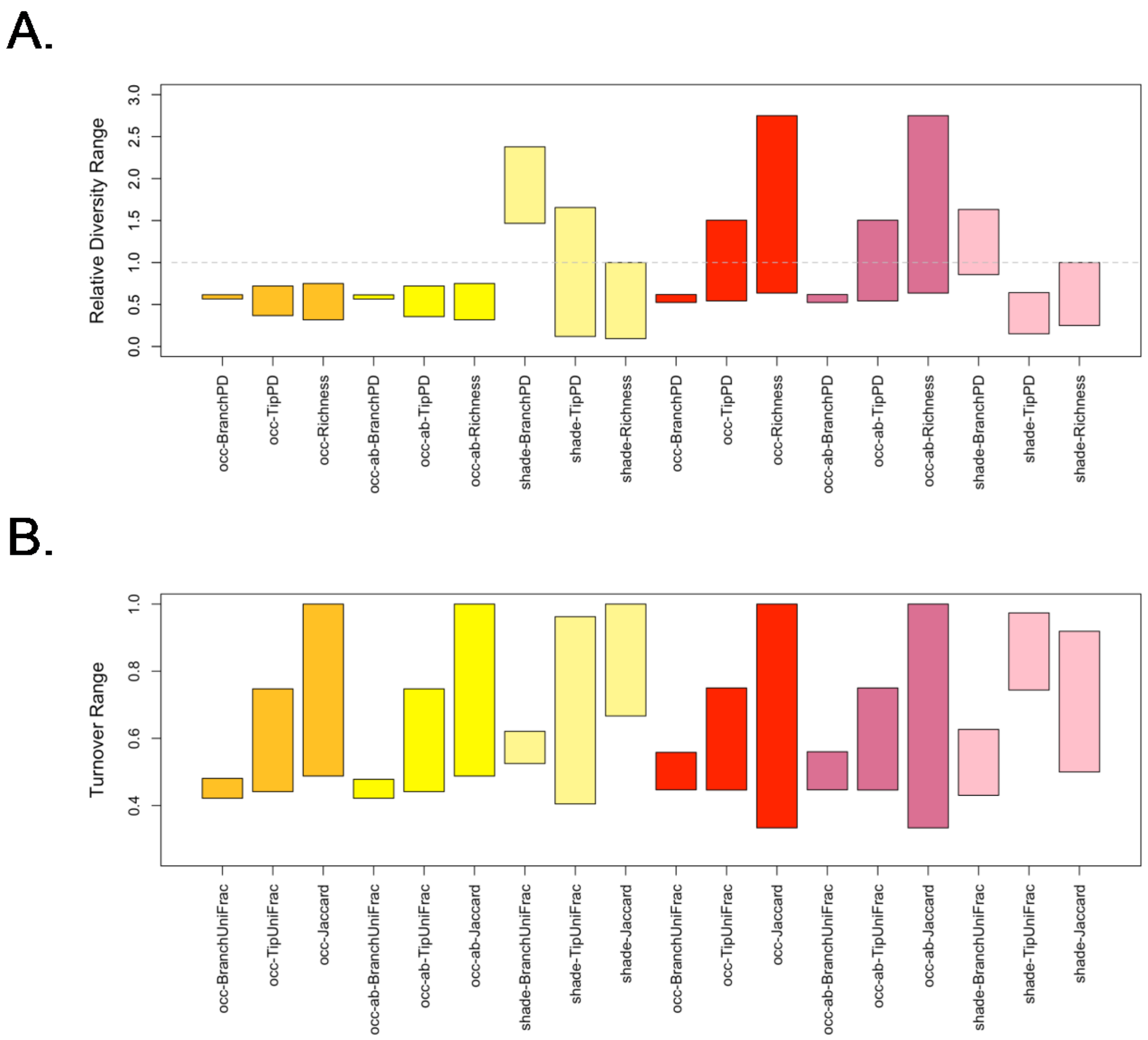


**Figure S5.1** Bar graphs showing minimum and maximum values of (A) alpha and (B) beta diversity across taxonomic rank for *Aspidoscelis arizonae* (gold, yellow, light yellow) and *A. marmoratus* (red, dark pink and light pink) relative to *A. neomexicanus*. In this figure, we use a 10% increase in Bray-Curtis/weighted UniFrac distance as our Shade and Stopnisek stopping criterion, as opposed to the default 2% used in the main text.
